# Phylogenetics, evolution and biogeography of four *Digitaria* food crop lineages across West Africa, India, and Europe

**DOI:** 10.1101/2025.03.27.645812

**Authors:** George P. Burton, Paolo Ceci, Lorna MacKinnon, Lizo E. Masters, Philippa Ryan, Colin G.N. Turnbull, Tiziana Ulian, Maria S. Vorontsova

## Abstract

**Background and Aims:** Millet crops in the grass genus *Digitaria* include white and black fonio (*D. exilis* and *D. iburua*), raishan (*D. compacta*) and Polish millet (*D. sanguinalis*), cultivated across West Africa, India, and Europe. Fonio and raishan crops are important to supporting food security and subsistence agricultural systems in rural communities, while *D. sanguinalis* is no longer cultivated. These crops are resilient to challenging climates. We aim to produce an integrated study of these crops: a phylogeny of the *Digitaria* genus including all four food species, to identify key crop wild relatives (CWRs); time-calibrated biogeographic analysis, to investigate the history and evolution of *Digitaria*; and morphological study to assess the transition between wild and domesticated species.

**Methods:** We use the Angiosperm 353 target-enrichment sequencing approach to produce maximum likelihood and coalescent model nuclear phylogenies for 46 *Digitaria* species, including ploidy estimations, and bayesian methods to propose an evolutionary and biogeographic history for the genus. Morphology of wild and cultivated species is investigated for spikelets and growth habits using microscopy and SEM imaging techniques.

**Key Results:** Four distinct lineages are proposed for the evolution of *Digitaria* crops, and close CWRs *D. fuscescens*, *D. atrofusca*, *D. setigera* and *D. radicosa*, and *D. ciliaris* are identified. South and eastern Africa is proposed as the origin of early *Digitaria* divergence, with crop lineages diverging from wild relatives around 2-6mya. Incomplete domestication traits are observed, including the loss of trichomes, but no significant difference in spikelet or abscission zone morphology.

**Conclusions:** *Digitaria* crops have important climate-resilient traits and hold strong potential for stabilising vulnerable food systems against the challenges of future climate change. The knowledge produced in this study about CWRs will be useful in improving crop traits through targeted breeding and indigenous crop conservation programmes.

## Introduction

Cereal crops provide the majority of the world’s food source, yet are likely to be disrupted heavily by future and present climate change (Reeves *et al*., 2016; Fatima *et al*., 2020). Many underutilised or ‘orphan crops’ are likely to be more tolerant to this change, and can provide resilience in vulnerable food systems that will otherwise be degraded (Ulian *et al*., 2020; Mabhaudhi *et al*., 2019; Santhoshini *et al*., 2025). A selection of these crops have been presented as Neglected and Underutilised Species (NUS; Ulian *et al*. 2020; Animasaun *et al*. 2023), which have experienced far less research and commercial development attention than major crops, despite their climate-resilient traits and importance to indigenous identity and rural communities. Many of these are millet grasses, including white (*Digitaria exilis* (Kippist) Stapf) and black (*Digitaria iburua* Stapf) fonio from West Africa. During intense periods of selection, major cereals have experienced a reduction in genetic diversity, which has also reduced their natural adaptability and trait plasticity. Under-improved crops (including fonio), and crop wild relatives (CWRs), are known to retain genes and allelic diversity that may have disappeared from the more heavily domesticated species which experience genetic bottlenecks, and hold the key to preventing diseases, increasing nutrition, and providing climate resilience in vulnerable areas (Warburton *et al*., 2006; Dempewolf *et al*., 2017; Kashyap *et al*., 2022; Hajjar and Hodgkin, 2007). The involvement of wild species in crop breeding programmes has been utilised extensively for rice (Stein *et al*., 2018; Zheng *et al*., 2024), wheat (Charmet, 2021), and maize (Warburton *et al*., 2017; Abdoul-Raouf *et al*., 2017). However, for the improved utilisation of orphan millets like fonio to be possible and effective, scientists first require a detailed understanding of the taxonomy, evolutionary history, and geographic niche of the study species. This knowledge of how these grasses have evolved and adapted to climatic conditions in the past, often indicates how and why they might respond to challenging future climates.

### *Digitaria* species uses and systematics

In the genus *Digitaria* Haller (Panicoideae: Poaceae) there are four crop species which are or have been incredibly important to agriculture systems and providing food for people: white fonio (*D. exilis*), black fonio (*D. iburua*), raishan (*D. compacta* (Roth) Veldkamp), and Polish millet (*D. sanguinalis* (L.) Scop.)(Cruz *et al*., 2016; Portères, 1955, 1957, 1976; Singh and Arora, 1972). The genus includes the cultivated pasture grass *Digitaria eriantha* Steud. (Pangola grass), common in South Africa, the Americas, and Southeast Asia (Parsons, 1972; Tikam et al., 2013). *Digitaria* grasses can also have negative economic impacts on agriculture, as the invasive, spreading habit of some wild species (including wild *D. sanguinalis, D. ciliaris* (Retz.) Koeler, and *D. humbertii* A.Camus) makes them difficult to manage in farm and horticultural settings (Jones *et al*., 2021; Touafchia *et al*., 2023; Randrianarimanana *et al*., 2024). Despite their importance, *Digitaria* grasses are challenging to identify to species level. Although there have been several genomic and phylogenetic studies of *Digitaria*, described below, there has been no integrated approach which considers phylogenetic, morphological, and biogeographical analysis of the group, also including both all 4 domesticated crop species, and close wild relatives. By producing this analysis, we can build knowledge of the close wild relatives of *Digitaria* crops and make connections to changes in past climates, and make it easier for future researchers to conduct research on this genus.

The genus *Digitaria* was first described by Haller (1768), and now contains around 250 species globally (POWO, 2025), occupying both temperate and tropical niches. Studies in *Digitaria* systematics include work by Stapf (1915), Henrard (1950), and Vega *et al*. (2009) who use morphological traits to construct a classification system for the group. Key traits that have been used to identify species include the shape and size of the upper glume, trichomes on spikelet veins, branching patterns of the inflorescence (technically a synflorescence (Kellogg, 2015), but referred to in this study as an inflorescence for ease of terminology), number of racemes, and number of spikelets in a group (binate = two spikelets, ternate = three or more). These characters are illustrated in Figure S1.

More recent studies in molecular phylogenetics have utilised the combination of ITS sequences and morphology (Lo Medico *et al*., 2017), plastid sequences (Morrone *et al*., 2012), and a mix of Sanger (ITS, *matK, rbcL, ndhF, atpB*, and *trnL-F*) and target-enrichment sequence capture of 353 genes (using the Angio353 bait kit) within a supertree matrix approach (Touafchia *et al*., 2023), to untangle the complex relationships between *Digitaria* grasses. Regional taxonomic treatments of *Digitaria* have been conducted in Pakistan and Central Asia, South East Asia, and Madagascar, by Gilani *et al*. (2003), Boonsuk *et al*. (2016) and MacKinnon *et al*. (in press) respectively. Minoji and Sakai (2024) in the assembly of a chromosome-level genome for the Asian species *Digitaria radicosa* (J.Presl) Miq., also included several of the major clades within a phylogenomic analysis. Throughout these studies several key patterns and traits have remained consistent: the topology includes several major clades distinguished by ternate vs. binate spikelet morphology, combined with the presence of spikelets with complex or absent vs. simple trichomes. The genus contains many polyploid species, and likely reflects a complicated history of introgression and hybridisation events, which makes genetic study of the group particularly difficult (Abrouk *et al*., 2020; Minoji and Sakai, 2024). The *Digitaria* clade has also given rise to other morphologically divergent grass genera including *Anthephora* Schreb. and *Chlorocalymma* Clayton (Touafchia *et al*., 2023).

### Food crops in *Digitaria*

White fonio (*Digitaria exilis*) is the most commonly cultivated crop in *Digitaria*, grown in West Africa, where it is especially important for both subsistence agriculture in rural areas (Cruz *et al*., 2016; Portères, 1976; Burton *et al*., 2024), and as an increasingly popular export crop (CBI, 2022; Tridge, 2023). It is recognised for its tolerance to drought and heat, high nutritional value, and socio-economic and cultural importance. However, it produced remarkable small grains (typically1-2mm in length), which pose post-harvest processing challenges due to the presence of a seed husk that is difficult to remove (Cruz *et al*., 2024; Animasaun *et al*., 2023; Portères, 1976).

The second most cultivated *Digitaria* species for food is black fonio (*Digitaria iburua*). This is restricted to small areas of Benin, Togo, and Nigeria, just below the Sahel region. There are few detailed studies available on this rare crop. Portères (1955) describes the likely domestication of *D. iburua* in the Aïr mountains of Niger before being transported south through to Nigeria, and then to northern Benin and Togo. It is relatively less popular than white fonio as it is reportedly even more difficult to de-husk, and in Togo it is instead used to make a beer called “tchapalo” (Cruz *et al*., 2016). White and black fonio plants are shown in Figure 1.

**Figure 1.**
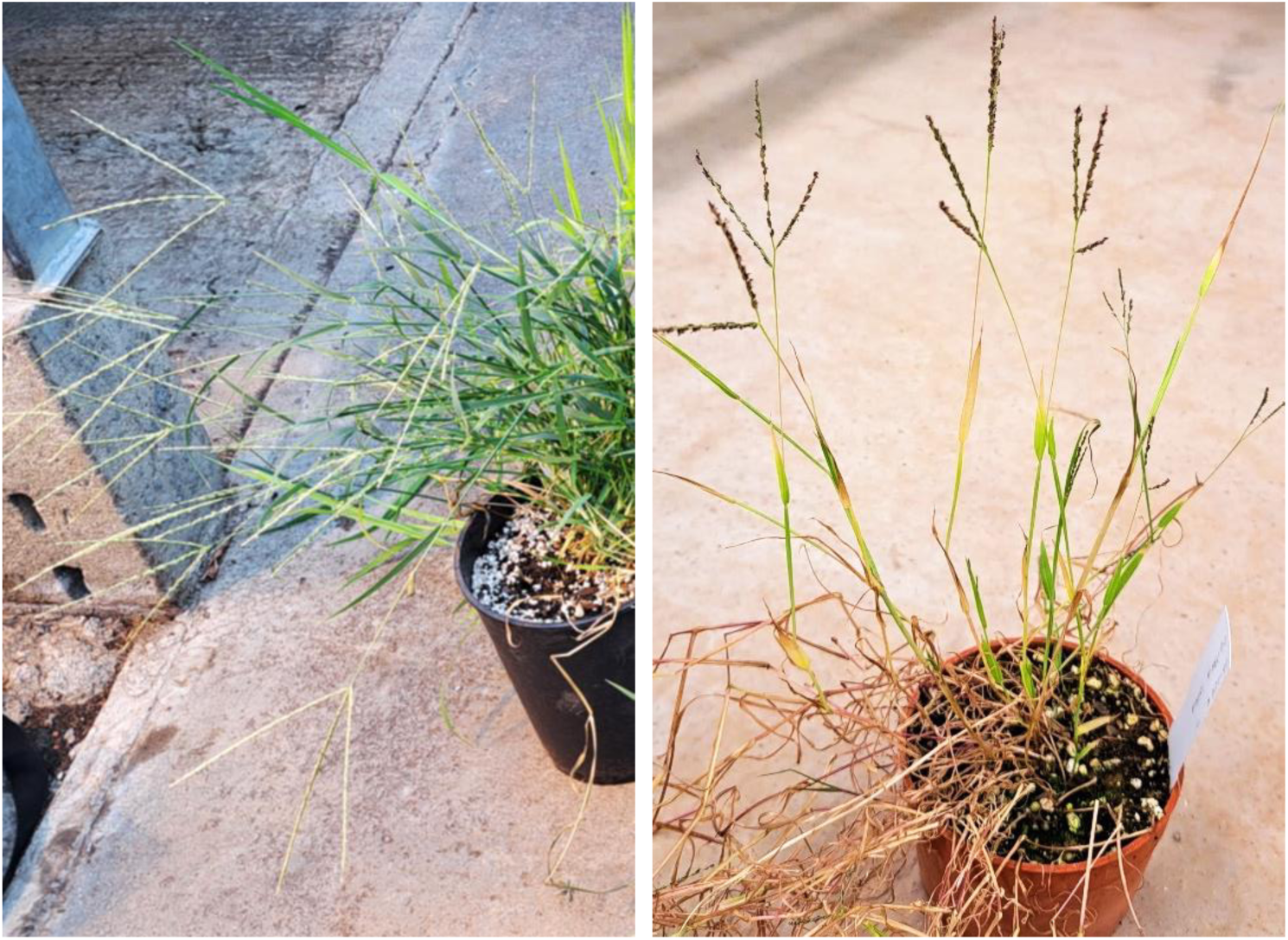
White (left) and black (right) fonio (*Digitaria exilis* and *D*. *iburua*), cultivated from seed originating in Guinea and Nigeria respectively, in the Jodrell Glasshouses, Kew. Photo credit to lead author.

A third *Digitaria* food crop, raishan (*Digitaria compacta*) is a millet cultivated in the Khasi Hills near Shillong (Meghalaya, India), a wet highland environment. Hooker (1857) describes the cultivation of what was earlier named as *Paspalum sanguinale* var. *commutatum* (auct. non Hook.f.) in nearby Assam and the Khasi Hills. Portères records that in the 1950s it was still under cultivation on about 100 acres of land in nearby Assam (Portères, 1957). The most detailed published source of information for this crop is a study by Singh and Arora (1972), which includes a theory of how the crop was selected and domesticated from wild *D. cruciata* (Nees ex Steud.) E.G.Camus & A.Camus, including photos of the crop’s long, unbranching racemes, providing details and evidence of its cultivation in tribal communities. This study makes no mention of cultivation in Assam, and since this study there have been no formal published reports of cultivation in Meghalaya. The name *Digitaria cruciata* var. *esculenta* Bor., previously used for the cultivated variety, is now an accepted synonym of *D. compacta* (POWO, 2025), a species with overlapping occurrence to wild *D. cruciata* across central and south-eastern Asia. A photo of raishan grown in the Khasi hills is reproduced from Singh and Arora (1972) in Figure S2.

Polish Millet (*Digitaria sanguinalis*) is thought to have been cultivated during the 1500-1600s in Eastern Europe, first intentionally cultivated in a monastery garden in Croatia or Albania, before moving north and expanding into Germany, Hungary, and later Poland and Ukraine (Portères, 1955 and 1957), before its likely decline in the mid to late 1800’s. Portères suggests that the first record of this cereal is from 1561, and that it was reported to still be cultivated in Poland in the 1890s. Netolitzky (1914) says that ‘blood millet’ was cultivated in Germany in the 16^th^ century. He found no difference between cultivated and wild grains. Henrard (1950) also mentions a specimen of the cultivated form of this crop (*D*. *sanguinalis* var. *esculenta* (Gaudin) Caldesi) at the Paris herbarium, which is described to have longer inflorescence racemes and a more robust, erect habit than common *D. sanguinalis*, however no specimens or detailed records could be found for this study.

### Fonio Genetic Studies

One of the earliest attempts to investigate the relationship between white and black fonio and their wild relatives using phylogenetic methods was by Hitu *et al*. (1997), who used Random Amplified Polymorphic DNA (RAPD) markers to support that the phylogenetically closest wild relative of white fonio is *D. longiflora* (Retz.) Pers., and that of black fonio is *D. ternata* (A.Rich.) Stapf. This relationship was tested and confirmed in later analyses by Adoukonou-Sagbadja *et al*. (2010), Nyam *et al*. (2017), and Ngom *et al*. (2017). Adoukonou-Sagbadja *et al*. (2010) supports a genetic affinity of over 92% between fonio crops and closest wild relatives, and a similar close ancestry between *D. sanguinalis* and the common weed *D. ciliaris* (Retz.) Koeler. The production of a chromosome-level annotated genome for *Digitaria exilis* by Abrouk et al., (2020) also confirmed the relationship between white fonio and *D. longiflora*, and shared useful insights into synteny shared with other cereal crops.

### Domestication of grass species

In the transition of a species from wild to a cultivated cereal crop, there is a domestication syndrome which involves phenotypic development based on traits which have been selected for by farmers (Meyer *et al*., 2012; Fuller *et al*., 2023). Through this form of ‘accelerated evolution’, wild plants experience selection through many generations of seed collection and re-sowing (Harlan *et al*., 1973). Estimates suggest that transition to early stages of domestication can take as long as 2000 years of active cultivation, but different crops display different thresholds of domestication phenotypes over time, dependent on plant pollination and reproductive strategies, and introgression, crossing, and cultivation methods (Brush, 1995; Allaby et al., 2017; Allaby et al., 2022; Glémin and Bataillon, 2009).

A key observable phenotype of domestication for many cereals and pulses is a decrease in seed shattering: mechanistic changes involving the devolution of an abscission zone occur in the inflorescence which prevents the grain from easily separating from the rest of the plant before it has reached full maturity (Yu and Kellogg, 2018; Yu *et al*., 2020). Plants with this trait increase their chances of being harvested and cultivated, unlike wild plants which benefit from dispersing their seeds (Woodhouse and Hufford, 2019). This is often accompanied by the loss or reduction of trichomes or awns on the spikelet - in wild plants the presence of trichomes allow the seed or spikelet to be dispersed by the wind and by animals, and help to establish the seed in the ground. Larger seed size is also a common sign of domestication, reflecting improved seed vigour for successful germination, especially for grasses in arid or difficult environments, and selection for higher yield and seed nutritional content (Fuller *et al*., 2023). The phenotypic signs for domestication between wild and cultivated species can be difficult to track due to variability of traits between and within species, especially in already difficult to identify and manage grasses such as *Digitaria*.

### Aims and Scope

The study of wild *Digitaria* grasses in relation to food crops including fonio has been overlooked, and understanding the evolution of the genus will provide valuable knowledge to future cultivation and biodiversity projects. This study aims to identify crop wild relatives within *Digitaria* beyond *D. longiflora* and *D. ternata*, and resolve when, where, and how separate crop lineages within *Digitaria* may have evolved.

We construct a well-resolved phylogeny of the genus using target-enrichment sequencing and both maximum likelihood and coalescent-based tree-building methods, with sampling taking into account geography, and focusing on clades containing crop domestications. We estimate the evolution and dispersal of *Digitaria* grasses from their early diverging species through to domesticated lineages, using both biogeographical and time-calibration methods, and record and discuss the development of distinct and shared morphologies across the group, including domestication traits. This provides an integrated history of the genus, involving the spread and growth of wild species as an important part of this crop history.

## Methods

### Species sampling and material

Ninety-six samples of 46 species were chosen for DNA extraction and sequencing based on the cladistics and monography presented by Henrard (1950) and Stapf (1915), Lo Medico *et al*. (2015), Vega *et al*. (2009), and Touafchia *et al*. (2023). To approach questions about the evolution of crop species, sample selection was targeted on close relationship and geographic affiliation with key crop species *Digitaria exilis*, *D. iburua*, *D. compacta*, and *D. sanguinalis*. *Urochloa ruziziensis* (R.Germ. & C.M.Evrard) Crins was chosen as the outgroup taxon, within Paniceae. Details of collections involved in the final trees are included in Table S1.

Samples were primarily collected from dried herbarium specimens available from the K herbarium, with few others from P herbarium (herbarium codes used from Index Herbariorum; Thiers, 2025), and others extracted from silica-dried material from recent fieldwork expeditions. Leaf and inflorescence material was sampled for DNA extraction. Specimens were identified and selected by comparison against Henrard (1950) and against the K herbarium collections, with a focus on type specimens. Type specimens of *Digitaria iburua* and *D. cruciata* var*. esculenta* (to represent *D. compacta*) were sampled at the K herbarium, from native cultivation regions in Nigeria and India, respectively. For material most likely to represent *D. sanguinalis* in its cultivation zone in Europe, a germplasm accession was acquired from the Leibniz Seedbank (IPK) in Germany, which was previously stored and grown annually during the summer in the Alter Botanischer Garten, Gottingen, described as a fast-growing invasive and a descendent of the cultivated variety. This material was grown and sequenced as part of the NERC Environmental Omics Facility (NEOF) de novo project NEOF1392 2021, using WGS (included in this study as sample GB999), and will be assembled and published as part of a future ongoing study.

Sixty-four target-capture DNA sequences and corresponding morphological traits were recovered and analysed for 46 species: 22 binate, 23 ternate, and one with solitary spikelets. Samples were most commonly taken from specimens collected in Nigeria (8), Madagascar (7), India (9), and Ghana (7). Of all species’ native distributions, 34 occur in Africa, 16 in Asia, 4 in the Americas, and 3 in Europe.

### DNA extraction and library preparation

DNA was extracted and isolated from samples using an adapted CTAB method (Doyle and Doyle, 1987), using CTAB, isopropanol and chloroform for purification, and magnetic beads for cleaning steps. Target capture sequencing was conducted using the Angiosperm353 target capture probe kit (Baker *et al*., 2021; Johnson *et al*., 2019). Libraries were prepared using indexes from the NEBNext Ultra II DNA library prep kit, and NEBNext Multiplex Oligos from Illumina. Pooled libraries were hybridised with baits from the Angiosperm353 probe kit for 24 hours at 65°C and 12 cycles of PCR. Samples were sent to Macrogen Inc. in Seoul, South Korea for sequencing using an Illumina HiSeqX platform, for 2x150bp paired-end reads. Tapestation (Agilent Technologies 4200 TapeStation) High Sensitivity D1000 ScreenTapes, gel electrophoresis and Quantus Fluorometer (Promega, USA) measurements were used for validating DNA integrity and quantity throughout the extraction and library preparation process.

### DNA sequence assembly

Raw data reads were trimmed of their adaptors and organised into paired-end and unpaired sequences, and filtered for quality using Trimmomatic 0.39 (Bolger *et al*., 2014). Loci assembly was completed using HybPiper 2.1.2 (Johnson *et al*., 2016) using a DNA target fasta file to capture reads from the Angiosperm 353 genes and WGS, using BLASTX v2.5.0 (Camacho *et al*., 2009). The gene coding regions were extracted and distributed into reads per 353 genes, per sample, using HybPiper. Average gene recovery statistics were provided by the ‘hybpiper_stats.py’ function in Hybpiper. Sequences were concatenated between genes, and aligned using MAFFT v7.476 (Katoh and Standley, 2013), then filtered to remove areas with <0.3 missing data within alignments using PhyUtility 2.2.6 (Smith & Dunn 2008).

### Phylogenetic tree-building

Cleaned gene alignments were 1) used to build a concatenated supermatrix using the -t1 flag ML method, with a GTR+G+I model, in IQTREE 2.1.2 (Minh *et al*., 2022), and 2) analysed with IQTREE to produce individual maximum likelihood gene trees based on BIC scores, low support values <0.3 were again collapsed using Newick Utilities 1.6 (Junier & Zdobnov, 2010), taxa with long branches removed using TreeShrink 1.3.9 (Mai and Mirarab, 2018), and finally a coalescent species tree was inferred using ASTRAL-III 5.7.7 (Zhang *et al*., 2018). Bootstrap values for ML trees were calculated using 1000 ultrafast replicates in the IQTREE concatenated tree, and LPP and quartet scores (QS) using the - t2 flag in ASTRAL-III for the coalescent species tree.

Visualisations of trees are produced in R environment version 4.4.0 (R Core Team, 2024), using the ggplot2 (Wickham, 2016), ggtree (Yu *et al*., 2017), ape (Paradis and Schliep, 2019), phytools (Revell, 2012), and treeio (Wang *et al*., 2020) packages.

A principal component analysis of sequence genetic distance was produced, using AMAS 1.0 (Borowiec, 2016) to concatenate gene alignments from MAFFT, and then a distance matrix of sequences was produced using the dist.alignment function in the seqinR (Charif and Lobry, 2007) to calculate a pairwise distance matrix from DNA sequence alignments.

### Time-scaled tree calibration and biogeography

Alignments were input to Sortadate (v. 2020-12-05; Smith *et al*., 2018), to rank gene clock-likeliness based on tip-to-root variation, bipartition, and treelengths, using the ‘gene-shopping’ method, to avoid computational issues caused by inputting large gene datasets in BEAST analysis. Alignments were concatenated and then split by gene partition using AMAS, to fill missing taxa per alignment. The top 15 ‘best’ genes were input as 15 separate partitions on BEAUti2 (https://www.beast2.org/beauti/).

An uncorrelated lognormal relaxed clock model was used to allow for independent evolutionary rates across taxa. A GTR site model (same as the ML tree) was used, with a gamma category of 4. Topology (the starting tree) was constrained to the rooted ML tree. A Yule model prior was set, with *Urochloa ruziziensis* set as an outgroup using a mean divergence time of 21mya (Hackel *et al*., 2019; Pessoa-Filho *et al*., 2018) and a sigma of 2. The MCMC run was set to 500,000,000 with a burnin of 50,000,000 (10%), sampling every 100,000. The .xml control file from BEAUti2 was then run using BEAST2 (Bouckaert, *et al*. 2014). Two independent chains were run and combined. The log file from these BEAST2 runs were analysed using Tracer (Rambaut *et al*., 2018), and tree files with an ESS >200 combined using LogCombiner (https://beast.community/logcombiner). A final MCC tree with mean node heights was produced using TreeAnnotator (https://beast.community/treeannotator).

The MCC tree from BEAST analysis and a set of binarised species distributions set into 5 main regions: Americas, West Africa, South and East Africa, Europe, and Asia, were using with the programme BioGeoBEARS (Matzke, 2013) in R to estimate and illustrate ancestral geographic ranges. Native species distribution data was collected from POWO, GBIF, and specimens at K herbarium. The models DEC and DIVALIKE (and +J) were used, and the best iterations were determined using LcL values.

### Ploidy Estimation

Ploidy level for samples was estimated using nQuire (v. 2018-05-05; Weiß *et al*., 2018). Sorted BAM files from the HybPiper gene extractions were input to nQuire, and analysed using the ‘lrdmodel’ function. Delta log-likelihood values were calculated and used to interpret ploidy (either diploid, triploid, or tetraploid and above). Reported ploidy levels for *Digitaria* species were accessed from the Chromosome Count Database (http://ccdb.tau.ac.il/).

### Morphological traits

Morphological traits were extracted from Plants of the World Online (POWO, https://powo.science.kew.org/), and GrassBase (Clayton *et al*., 2016), and verified against specimens and type collections at K herbarium. Spikelet traits were observed using dissection microscopes. Traits collected included growth form (mat forming, rhizomes or stolons), growth habit (erect, spreading), whether spikelets were binate or ternate, presence and type of trichomes present on spikelets (glabrous, complex or simple trichomes), number of racemes, length of racemes, and length and width of spikelets.

### Imaging

Images of whole plants including inflorescences and roots were taken using an Epson 10000XL scanner at 800dpi and processed using Adobe Photoshop. Spikelets were imaged using a Leica Z-stacking microscope. *Digitaria* spikelets were drawn to 1.25mm scale at 24x magnification by Lucy T. Smith.

SEM was performed using two methods: the first using platinum coating at the Jodrell Laboratory in Kew Gardens, and the second using VP-SEM with uncoated spikelets at the Natural History Museum, London. Spikelets were samples from herbarium sheets at K. For SEM the samples were placed on metal stubs and coated with platinum, using a quorum-Q150t es Series sputter-coater. A Hitachi 8230 with a 5kV electron beam was used to process images. For VP-SEM, uncoated spikelets were placed on metal stubs, and observed using a FEI Quanta 650 FEG scanning electron microscope.

## Results

### DNA Sequence Recovery

DNA recovery statistics from the 353 target genes resulted in capturing an average of 345 reads, 309 mapped reads, 305 on target reads, 287 genes mapped, and 244 genes with contigs, per sample. Full recovery statistics from HybPiper are provided in Table S2. Morphological traits, reported ploidy, and clade group numbers of key cultivated and wild species are shown in Table 1, and mapped across phylogenetic trees in Figures S3 and S4.

**Table 1.** Morphological traits, distribution, and reported ploidy of *Digitaria* crop species and CWRs discussed in text. Distribution is given as native and non-native range, from POWO (https://powo.science.kew.org/). Ploidy is indicated as 2x (diploid), 4x (tetraploid), and 6x (hexaploid), with references.

| Species | Clade | Distribution | Role | Spikelet | Lower Glume | Upper Glume | Lower Lemma | Ploidy | Reference |
| --- | --- | --- | --- | --- | --- | --- | --- | --- | --- |
| *D. exilis* | 2 | W. Africa | Crop | Ternate | Absent | Full spikelet, glabrous | Glabrous | 4x | Adoukonou-Sagbadja et al., 2007, Abrouk et al. 2020 |
| *D. longiflora* | 2 | Global | Weed | Ternate | Absent | Full spikelet, trichomes | Trichomes | 2x, 4x | Hoshino and Davidse 1988, Adoukonou-Sagbadja et al., 2007 |
| *D. iburua* | 3 | W. Africa | Crop | Ternate | Absent | 1/2 spikelet, glabrous | Glabrous | 4x | Adoukonou-Sagbadja et al., 2007 |
| *D. ternata* | 3 | Global | Weed | Ternate | Absent | 1/2-2/3 spikelet, trichomes | Trichomes | 4x | Hoshino and Davidse 1988, Adoukonou-Sagbadja et al., 2007 |
| *D. compacta* | 5 | Asia | Crop | Binata | Small | 1/3 spikelet, trichomes | Glabrous | 2x, 4x | Mehra, P. N. 1982; Kumar and Subramaniam 1987; Löve 1985 |
| *D. cruciata* | 5 | Asia | Weed | Binata | Small | 1/3 spikelet, trichomes | Glabrous | 2x, 4x | Mehra, P. N. 1982, Koul & Gohil 1991 |
| *D. ciliaris* | 6 | Global | Weed | Binata | Small | 1/2 spikelet, trichomes | Trichomes | 4x, 6x | Carnahan and Hill, 1961, Trouin, 1972 |
| *D. sanguinalis* | 6 | Global | Weed, Crop | Binata | Small | 1/3 spikelet, trichomes | Trichomes | 4x, 6x | Šmarda et al., 2014, Xue et al. 1992, Bennet et al., 1998 |

### Ploidy Estimation

Twenty-eight samples were estimated to be diploid, 22 triploid, and 13 tetraploid or above, shown in Figure S4. Tetraploid or above samples include *Digitaria iburua*, *D. exilis*, *D. sanguinalis*, and *D. monodactyla* (Nees) Stapf., *D. gayana* (Kunth) A.Chev., and *D. leptorhachis* (Pilg.) Stapf. Diploid samples include *D. abyssinica*, *D. cruciata*, *D. compacta*, *D. ternata*, and *D. longiflora*. Samples of *D. ciliaris* and *D. sanguinalis* were estimated as both triploid and tetraploid.

### *Digitaria* species topology

There is similar topology for *Digitaria* species presented by the ML and coalescent method trees produced by IQTREE and ASTRAL-III respectively, shown in Figure 2. In both trees, there is a split into two large clades: clades 1-3 with almost exclusively ternate species (*D. monodactyla* in clade 1 being the only binate species), and clades 4-6 with exclusively binate species. Species inflorescence morphology for each food crop clade is shown in Figure 3.

**Figure 2.**
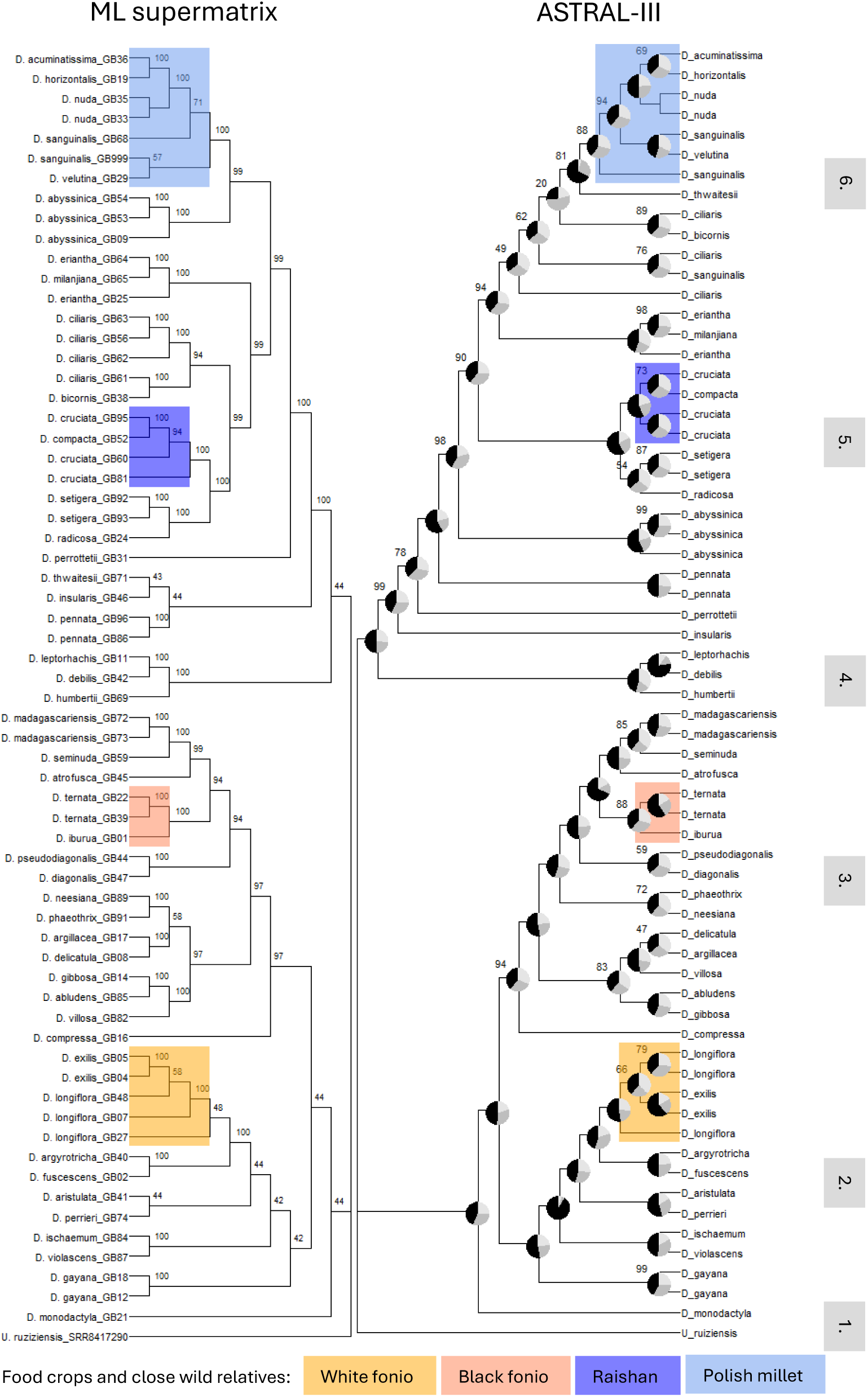
Maximum likelihood supermatrix (left) and ASTRAL-III coalescent (right) phylogenetic trees of *Digitaria* species. Bootstrap scores are displayed on the ML tree, and Quartet Scores and pies showing primary (black), second (medium grey) and third (light grey) gene tree topology agreement. Species with cultivated food species and close wild relatives are highlighted: fonio species in orange, and raishan and Polish millet species in blue. Clades are numbered 1-6 for discussion: 2-3 are ternate species, and 4-6 are binate.

**Figure 3.**
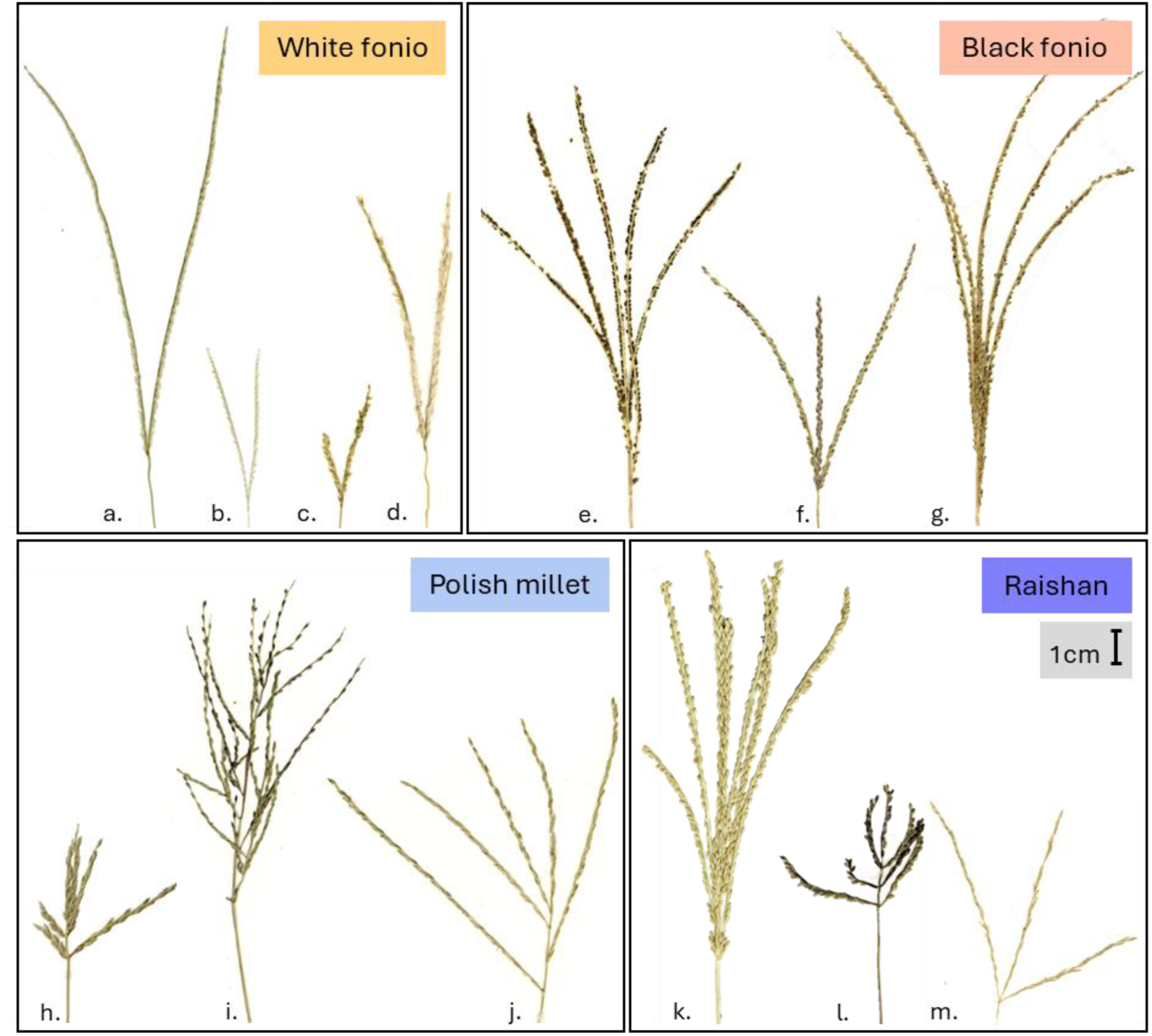
Photo scans of *Digitaria* inflorescences from herbarium specimens, representing the 4 main cultivated food crops, and closely related wild relatives. Species shown are: a. *Digitaria exilis* (DP 1001), b. *D. longiflora* (R.A. Farrow 112a), c. *D. fuscescens* (J.B. Hall 3304), d. *D. argyrotricha* (Greenway 10819), e. *D. iburua* (DP 1010a), f. *D. ternata* (Tuley 1547), g. *D. atrofusca* (A. Falannigui 25), h. *D. sanguinalis* (Stewart 26651), i. *D. velutina* (Hatch 4296), j. *D. ciliaris* (Gledhill 200), k. *D. compacta* (Hooker & Thomson s.n.), l. *D. cruciata* (Stainton, Sykes & Williams 3989), and m. *D. radicosa* (Vorontsova 2116). Scale bar shown to 1cm, all images to same scale.

#### Clade 1 – Digitaria monodactyla

*Digitaria monodactyla* is the first diverging species in the wider ternate group, though it is itself binate. The location of this primary divergence is the same on both trees with high ASTRAL-III support values: >99%, and gene tree congruence for first topology.

#### Clades 2 and 3 – White and black fonio and relatives

Within clades 2 and 3 the two different fonio crop species emerge into separate well-supported clades in both trees: clade 2 with white fonio (*D. exilis*) and relatives *D. longiflora, D. argyrotricha* (Andersson) Chiov., and *D. fuscescens* (J.Presl) Henrard (this sister clade supported by 100% ML bootstrap and QS), and clade 3 with black fonio (*D. iburua*) and relatives *D. ternata*, *D. atrofusca* (Hack.) A.Camus, and *D. madagascariensis* Bosser (sister clade supported at 94% ML and 100% QS). On the ASTRAL-III tree, low gene tree congruence in clade 2 separates multiple samples of *D. longiflora* and *D. exilis* (58% ML, 66% QS). One *D. longiflora* sample from Tanzania (GB27) is observed to be outside of the clade containing West African fonio and other *D. longiflora* samples (48% ML, 100% QS). Support for the separation between *D. iburua* and *D. ternata* in clade 3 is strong at 100% ML, and 88% QS.

#### Clade 4 – Digitaria leptorhachis, D. debilis and D. humbertii

A clade with *D. leptorhachis*, *D. debilis* (Desf.) Willd. and *D. humbertii* A.Camus contains the first diverging species among the binate species (not including *D. monodactyla*). This is congruent in both trees, with 100% ML and QS support, and especially strong gene tree agreement in the ASTRAL tree for the relationship between *D. debilis* and D*. leptorhachis*.

#### Clade 5 – Raishan and relatives

Both trees place the Indian crop raishan (*D. compacta*) in a well-resolved clade with relatives *D. cruciata*, *D. setigera* Roth. and *D. radicosa,* with identical topology between trees, and high ASTRAL-III support for the difference between *D. cruciata* samples and relatives *D. setigera* and *D. radicosa* (100%) and for the whole clade (90%). This is similarly high in the ML tree, with 94-100% support values for the whole clade and within.

#### Clade 6 – Polish millet, *D. ciliaris*, and relatives

The placement of *D. sanguinalis* samples is poorly supported in the ML tree (57-71%), but more confidently in ASTRAL-III (84-94%), where two samples from India (GB68) and Germany (GB999) appear more closely related to *D. nuda* Schumach. and *D. velutina* (Forssk.) P.Beauv., respectively.

The most discordant relationship between trees is the placement of species *D. abyssinica*, *D. eriantha*, and *D. ciliaris*. While the ML tree presents clade 5 and *D. ciliaris* as well-supported sister groups (99%), with samples of *D. ciliaris* from India, Brazil and Africa forming a single clade in the ML tree (94%), the ASTRAL-III tree splits the *D. ciliaris* samples with low support (20-62%), placing them between the *D. eriantha* and *D. sanguinalis* clades. *Digitaria eriantha* itself is well-supported as a clade with *D. milanjiana* (Rendle) Stapf. at 99-100% ML, and 94-100% QS, and appears to be closely related to *D. ciliaris* in both trees. A clade of *D. abyssinica* contains samples from across Africa and South America, all supported in the same clade together in both trees (99-100%), however it appears to be more closely related to *D. sanguinalis* in the ML tree (99%), while its placement outside of both *D. compacta* and *D. sanguinalis* clades in the ASTRAL-III tree is similarly well supported (98%).

#### Genetic distance matrix

A PCA of genetic distance between concatenated sequence alignments is shown in Figure 4 – the species *D. monodactyla*, *D. debilis*, *D. humbertii*, and *D. leptorhachis* appear to be the closest to the outgroup *Urochloa ruziziensis*, and the most separate and distant from either major binate or ternate clades. These distinct divergences are well-supported in both ML and ASTRAL-III trees above, which share the same main topologies in gene congruence.

**Figure 4.**
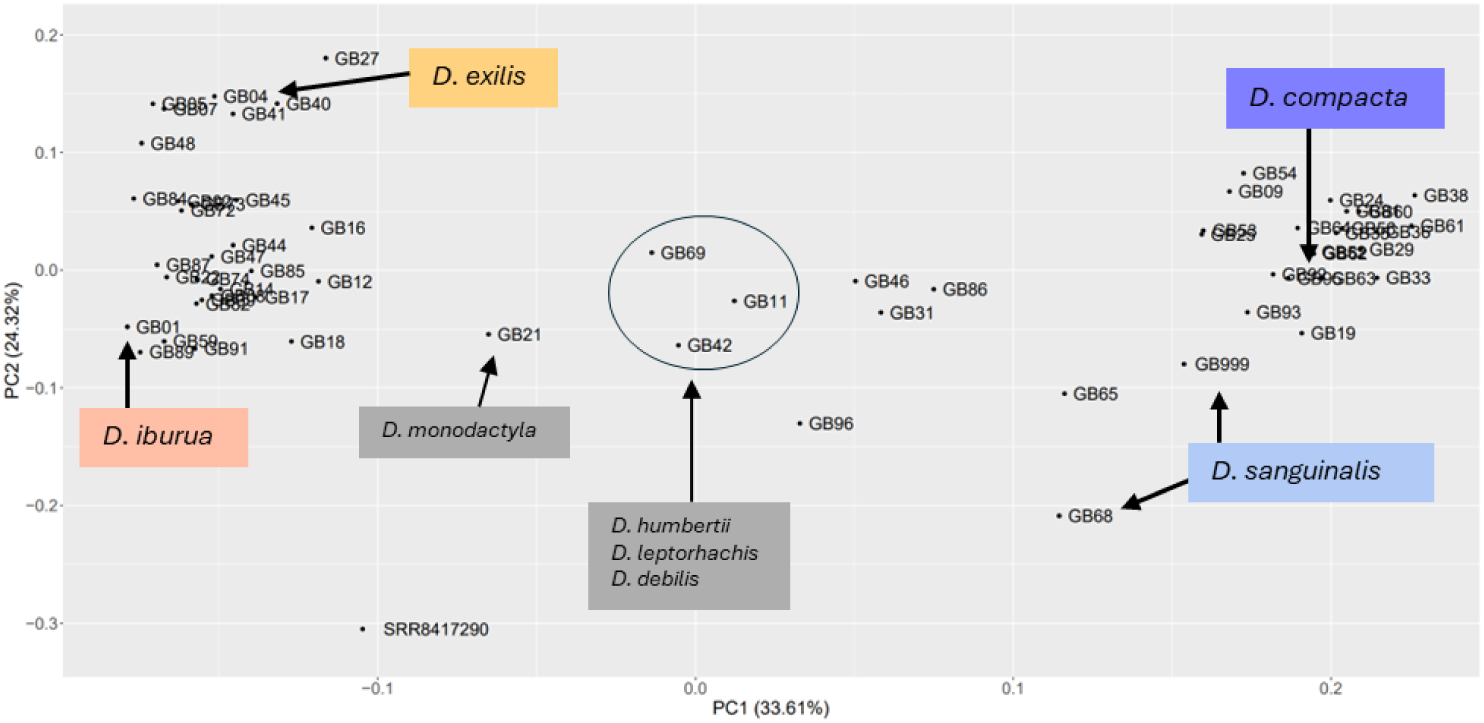
PCA plot of genetic distance between *Digitaria* sample alignments. Colours correspond to crop clades shown in Figure 2.4, key species indicated in labels, including early diverging *Digitaria* ternate species *D. monodactyla*, and binate group circled. Sample SRR8417290 is the outgroup species *Urochloa ruziziensis*.

### Evolution and biogeography

The results of the BEAST2 Bayesian analysis are shown in **Figure** 5. The earliest diverging extant *Digitaria* are African natives *D. monodactyla*, *D. compressa*, *D. gayana* from mostly ternate clades 1-3, and *D. debilis, D. leptorhachis,* and *D. humbertii* in binate clade 4. Both *D. exilis* and *D. iburua* are estimated to diverge from wild relatives around 2-3mya. *D. sanguinalis* diverges from *D. velutina* (north Africa) around 2.7mya, though an earlier divergence into India is dated at 6.22mya. The specimen of *D. compacta*, cultivated in India, diverges from wild *D. cruciata* more recently at 1.33mya, and both species from the clade containing *D. radicosa* and *D. setigera* around 6mya.

**Figure 5.**
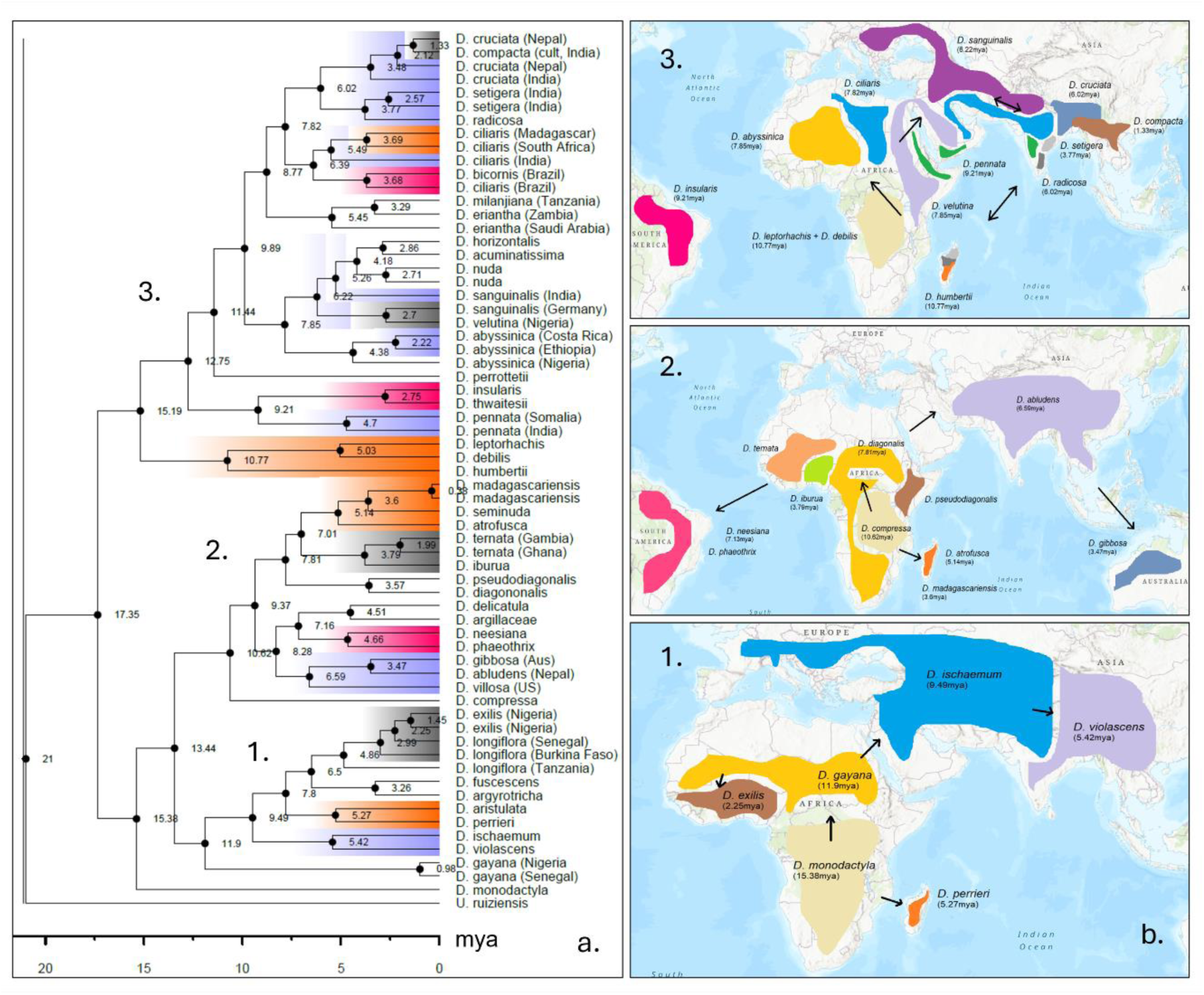
Bayesian (BEAST2) time-calibrated phylogeny of *Digitaria* species, and likely dispersal routes. **a.** Maximum clade credibility (MCC) tree, where clades are coloured by dispersals to Madagascar (orange) and the Americas (pink). Cultivated food species and relatives are highlighted in grey. **b.** Likely routes of dispersal, corresponding to clades numbered in MCC tree. Maps are based on overlapping native extant species distributions, from POWO (https://powo.science.kew.org/).

Biogeographical analysis by BioGeoBEARS is shown in Figure S5, which supports an ancestor of *Digitaria* in south-eastern Africa (DEC model LnL= -167.38, DEC+J Lnl= - 162.34). This corroborates the BEAST results where the earliest diverging *Digitaria* (including *D. monodactyla*, *D. gayana*, *D. compressa*, *D. leptorhachis*, and *D. debilis*) all have native distributions which overlap in south and eastern Africa.

### Domestication

Illustrations of *Digitaria* crop and wild species spikelets are shown in **Figure** 6. There appears to be a loss of trichomes for cultivated fonio species, though it is not fully complete. *Digitaria longiflora* has trichomes present on the spikelet and pedicel which are not present on *D. exilis*. Trichomes are present on the spikelet and pedicel of *D. ternata*, and on *D. iburua* there are no trichomes on the spikelet, but they are still present on the pedicel. Trichomes are present on the pedicel of both *D. compacta* and *D. cruciata*, and only slightly discernible on some specimens on the spikelet. On *D. sanguinalis* complex trichomes are very noticeable across the whole spikelet and pedicel. Close relatives *D. ciliaris,* and *D. velutina* (not pictured), are similarly pubescent.

**Figure 6.**
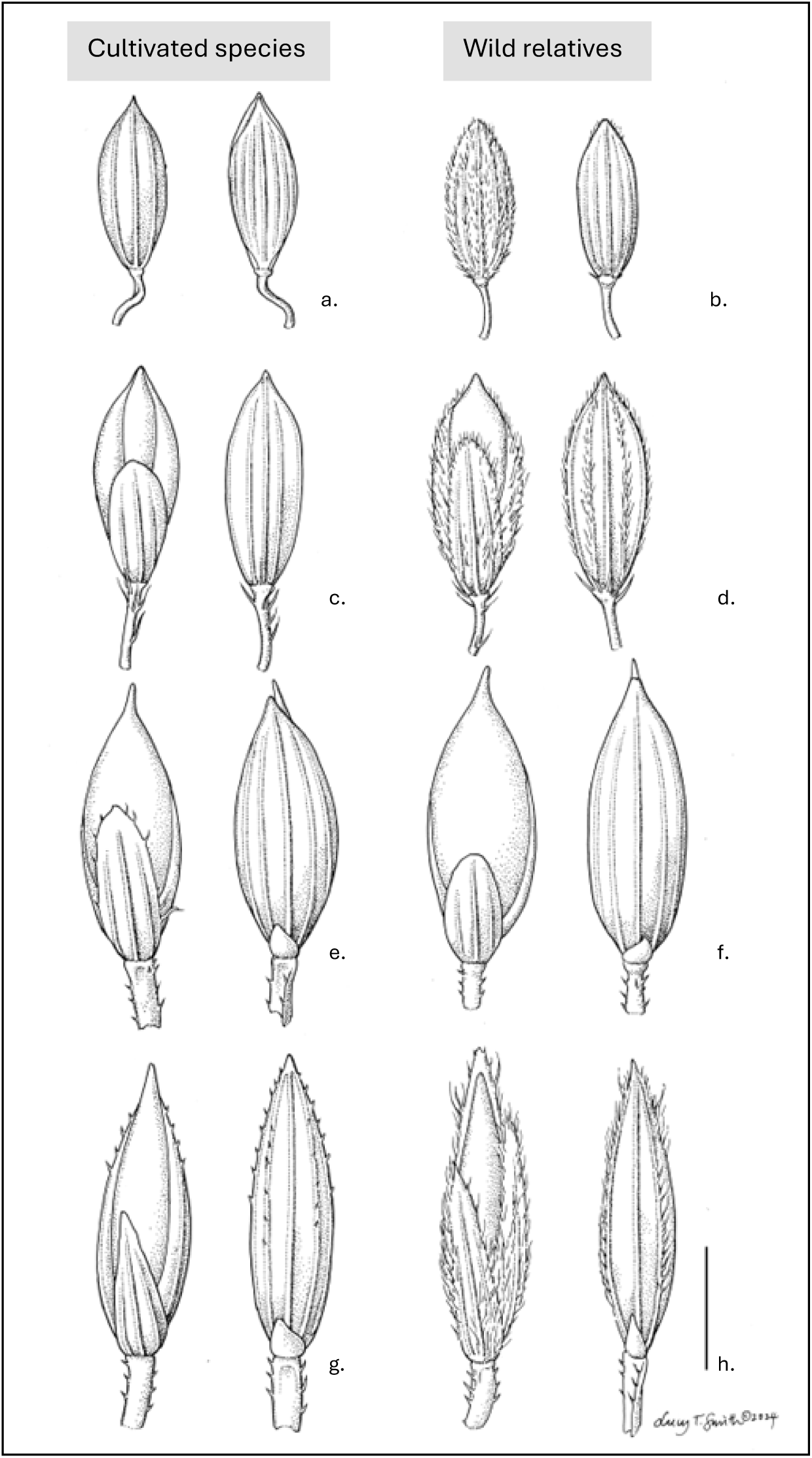
Illustrations of *Digitaria* spikelets, representing the 4 main cultivated food crops, and CWRs. Species are labelled from a-h: a. *Digitaria exilis* (Clarke s.n), b. *D. longiflora* (), c. *D. iburua* (Lamb 54), d. *D. ternata* (Schimper 76), e. *D. compacta* (Hooker & Thomson s.n.), f. *D. cruciata* (Royle s.n), g. *D. sanguinalis* (Stewart 26651), h. *D. ciliaris* (Lindeman & Haas 459). Drawn by Lucy Smith at Kew Gardens. Scale bar = 1.25mm at x24 magnification.

Regarding size of spikelet, from average measurements provided by GrassBase (Clayton et al., 2016), *D. exilis* has an average spikelet length of 1.7-2mm versus 1.2-2mm for *D. longiflora*, and *D. iburua* has ca. 2mm, versus 1.8-2.7mm in *D. ternata*. Although there are very few samples of cultivated *D. compacta* available, the average spikelet length is recorded as ca. 2mm, versus 2-3.5mm in *D. cruciata*, and 2.5-3mm in *D. radicosa*. *D. sanguinalis* has an average of 2.5-3.5mm, which is smaller than 1.5-2.1mm in *D. velutina*, but similar to 2.5-4mm for *D. ciliaris*. Visually, the crop species do not appear any larger than wild species, thorough future morphometric studies should be conducted on a wide range of material to confirm this.

The abscission zone for *Digitaria* species are shown in **Figure** 7 (cultivated and CWRs) and Figure S6 (wild species). The abscission zone is evident below the whole spikelet, shown by comparison of complete spikelet and removal, where an empty pedicel is shown. On the abscission zone itself, a layer of specialised cell structures appear to be arranged horizontally in squares, perpendicular to the vertical, long striations in the pedicel, likely representing lignification patterns. The same abscission zone cell patterns are seen in both wild *D. cruciata* and cultivated *D. compacta*, as is the ‘clean’ line pattern in both *D. longiflora* and *D. exilis*, and in *D. iburua* and *D. ternata*. The ‘horizontal cell’ pattern is only observed for binate species in *D. sanguinalis*, though this may be due to inconsistency of age between specimens. There appears to be no easily noticeable or consistent phenotypic difference between the wild and cultivated species. In Figure S6 an array of wild *Digitaria* species are shown for comparison, presenting the concave and convex sides of the pedicel base and abscission zone layer, which follows the same patterns as seen in cultivated and CWR species.

**Figure 7.**
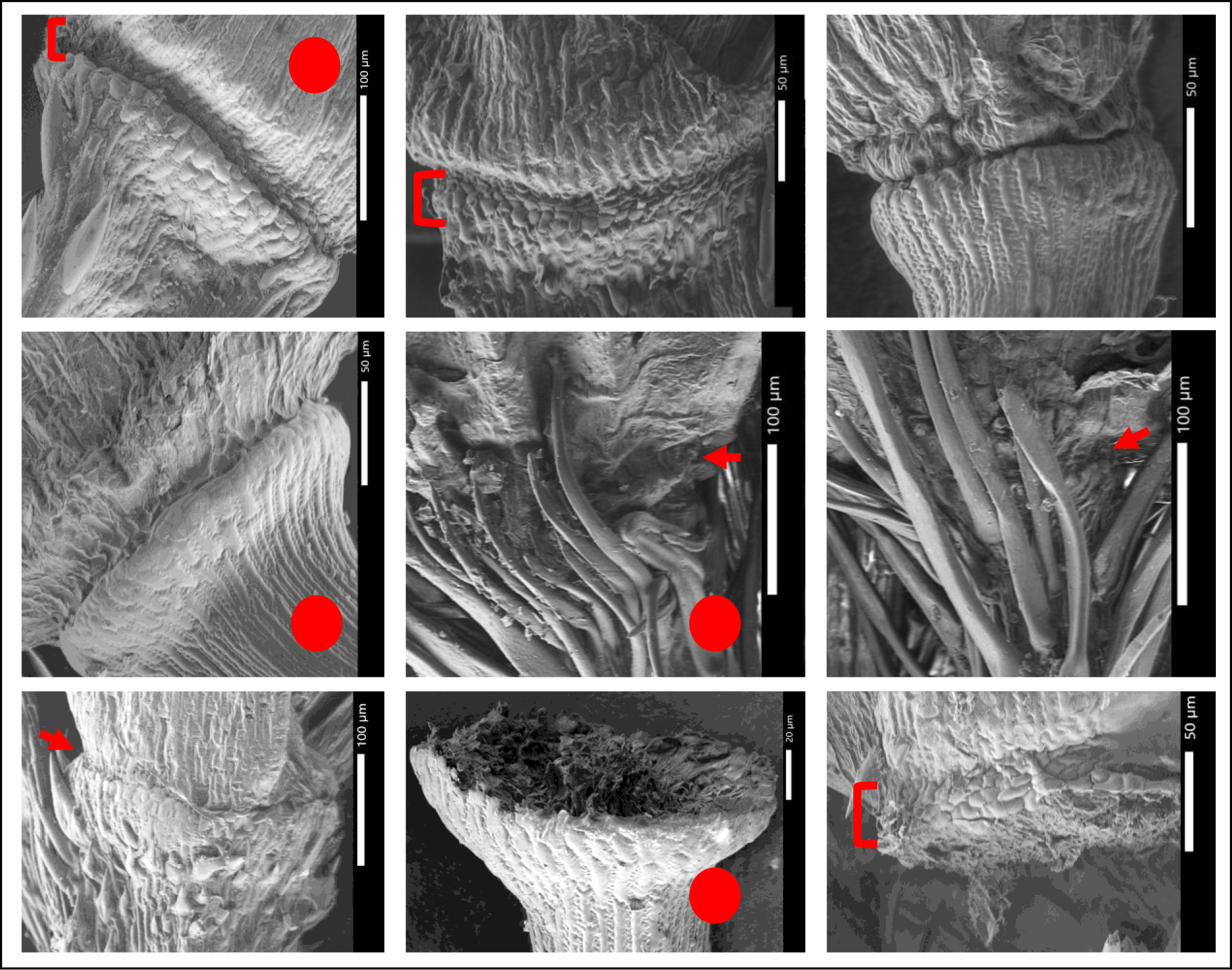
Scanning Electron Microscope photos of *Digitaria* abscission zones, cultivated and close wild relatives, captured at Kew Gardens and the Natural History Museum (UK). Red circles indicate cultivated species. Arrows and brackets indicate abscission zone area. Species from top left to bottom right are *D. compacta* (Hooker & Thompson s.n.), *D. cruciata* (Royle s.n)*, D. longiflora* (Merklinger 65), *D. exilis* (Burton 15), *D. iburua* (Lamb 54, type), *D. ternata* (Hall 636), *D. sanguinalis* (Santos 1844) *D. exilis* (Thomas 1485), and *D. sanguinalis* (Peterson 23883).

## Discussion

### *Digitaria* species topology

The structure of the nuclear trees and general clade topologies are similar to those of the nuclear trees presented by Touafchia *et al*. (2023), and Minoji and Sakai (2024). However, these trees are the first presentation of *Digitaria* using a high number of low-copy target nuclear genes, a wide range of species sampling both geographically and across all major clades, and including all 4 known species domesticated for food cultivation, and their close wild relatives. Our phylogenetic arrangement splits the species cleanly into binate and ternate species clades, contrary to the arrangement of species in a study by Touafchia *et al*. (2023), where ternate species are nested within two larger binate clades, though this may be affected by differences in sampling. The species *D. gayana*, *D. humbertii*, and *D. debilis* are also similarly early diverging in both studies.

#### Clade 2 – White fonio and relatives

The closest relative of *D. exilis* was again found to be *D. longiflora*, following previous studies including Adoukonou-Sagbadja *et al*. (2010) and Abrouk *et al*. (2020). A sample of *D. longiflora* from Tanzania, the only one sampled from outside of West Africa, is also seen to be less closely related to *D. exilis* in both trees, suggesting possible recent or ongoing introgressions between these two species in the range of fonio cultivation. The two species are similar in morphology and often grown in close proximity in cultivated fields across West Africa (Burton *et al*., 2024), though *D. exilis* has lost its trichomes, has increased raceme number and length, and abandoned the rhizomatous growth habit. The next nearest set of closely related species are *D. fuscescens* and *D. argyrotricha*, the former which is native to West Africa, and the latter to the coast of East Africa. All 4 species have maintained a strong ‘v-shaped’ finger inflorescence shape with no or rare auxiliary branching (see **Figure** 2). *D. exilis* is the only species which commonly has more than 2 racemes, though this varies between landraces, and the racemes are usually still connected at a single node. The morphology of *D. fuscescens* is similar to *D. longiflora* in the short racemes, and has similar ‘weedy’ invasive traits, such as a spreading habit, stoloniferous roots, and small inflorescences. *D. argyrotricha* has longer racemes with larger spikelets, and these are covered with a dense coat of trichomes. *D. gayana* is the earliest divergent ancestor in this lineage, an African native which varies between a ‘v-shaped’ and a sometimes more branched inflorescence, and spikelets also with dense trichomes.

There is clear morphological uniformity between the later-diverging 4 species, which appears to have been conserved over the course of around 7mya in Africa. White fonio has not become completely unrecognisable from its close relatives: its racemes are longer, but it still experiences lodging and shattering. Its lineage in virile, invasive, and drought tolerant African grasses is likely to afford significant unexploited potential, which in the future through breeding programmes and targeted selection may dramatically improve the food security of the rural communities that rely on it, otherwise suffering from a noticeably ‘incomplete’ domestication.

Analysis suggests that samples of the two fonio crops *D. exilis* and *D. iburua* are likely to be tetraploid (2n=36), corroborating previous study by Adoukonou-Sagbadja *et al*. (2007). However, the close relatives *D. longiflora* and *D. ternata* were estimated to be diploid – contrary to previous studies which suggest that both are tetraploid (Abrouk *et al*., 2020, Adoukonou-Sagbadja *et al*., 2007). Only one diploid accession of *D. longiflora* (2n=18) has been reported, by Hoshino and Davidse (1988). Close relative *D. fuscesens* is estimated here as tetraploid, and has been reported as both diploid and tetraploid in Gould and Soderstrom (1967 and 1974).

#### Clade 3 – Black fonio and relatives

The close relatedness of *D. iburua* and *D. ternata* has been reported since Hilu *et al*. (1997), and is supported again here. A sister clade containing the common African weed *D. atrofusca* is also presented. All three species share a common morphology: often multiple secondary branches from the main rachis, and distinct black upper florets – also shared with their earliest diverging species in the clade, the African *D. compressa*. Between the wild species and *D. iburua* we observe the development of thick culms, robust racemes, and the loss of spikelet trichomes. *D. ternata* and *D. atrofusca* overlap with *D. iburua* in its restricted cultivation area in Nigeria, and also both occur across West and south-eastern Africa. It is likely from this, their shared morphologies and phylogenetic similarity, that there were several speciation events within this cluster of species in West Africa as early as 8mya.

The documentation of black fonio cultivation and subsequent research on its traits and genetic history is particularly scarce, and recent sequencing and breeding projects aimed at improving knowledge about *D. iburua* (unpublished) often rely on germplasm provided by seedbanks in the US, Togo, Benin, and Nigeria. However, due to low germplasm quantities stored at these seedbanks, very limited germplasm could be obtained for this and other related studies, from a single accession collected in the 1960s. Authors were fortunately able to collaborate with Dr. Abubakar Bello and Muawiyya from Umaru Musa Yar’adua University, Nigeria, and early surveys from their work confirm that black fonio is still being cultivated in the Jos Plateau region. The scarcity of available germplasm material for black fonio is concerning, and more collections should be made to enable future research and conservation of an indigenous socio-economically important and arid-tolerant crop. Future research with this crop will benefit from including material not only from *D. ternata* but also *D. atrofusca* into their studies. Although *D. iburua* also suffers from apparent incomplete domestication in its small shattering grain, its robust growth habit and multiple racemes make it maybe even more attractive than *D. exilis* for breeding programmes, if it can be conserved, and promoted alongside improved access to de-husking and processing machinery.

#### Clade 5 – Raishan and relatives

The cultivated *D. compacta* is confirmed to be closely related to the wild *D. cruciata*, with a close sister clade including *D. radicosa* and *D. setigera*, all Asian grasses. Morphology-based cladistics of *Digitaria* from Pakistan and Central Asia in a study by Gilani *et al*. (2003) corroborate these findings, predicting a close relationship between *D. cruciata* and *D. sanguinalis* first, then followed by *D. ciliaris*, *D. setigera* and *D. radicosa.* There is a stark development from wild *D. cruciata* to the cultivated *D. compacta*: though they both have the same inflorescence structure with opposite secondary branchings from the same rachis node (in a ‘star’ shape), the cultivated form has longer racemes with more grains, and a more robust root to culm growth form. The inflorescence structure of sister clade *D. setigera* and *D. radicosa* are also similar in their parallel inflorescence branchings. Although all three species are common from Afghanistan eastward, *D. radicosa* and *D. setigera* also occur in Madagascar and other areas of south-eastern Africa – across which dispersal events may have taken place.

Raishan has long, numerous racemes, and spikelets with short trichomes, upper glumes to around ½ the length of the spikelet, all of which are reminiscent of both *D. sanguinalis* and *D. ciliaris*. Photos of raishan in the study by Singh and Aurora (1972; Figure S2) show it to have an impressive height, taller than either fonio species. The robust root to culm growth form is also similar to the form of *D. iburua*. The recently published chromosome-level analysis of close relative *D. radicosa* in Minoji & Sakai (2024) included an analysis of synteny between *D. radicosa* and other crops, finding strongly similar syntenic blocks between *D. radicosa* and *D. exilis*, and with rice (*Oryza sativa*), including a diverse range of NLR (resistance gene analog RGA5) genes to protect from blast fungi. *Digitaria radicosa* is likely to be a useful CWR species in future research analysing the genetic evolution and domestication of the crop raishan.

Both *D. cruciata* and *D. compacta* are estimated to be diploid in this study, and have been previously reported as diploid and tetraploid (Kumar and Subramaniam, 1987; Löve, 1985; Mehra, 1982; Koul and Gohil, 1991). Close relative *D. radicosa* is estimated to be tetraploid in this study, but was found to be exclusively diploid in a chromosome-level genome assembly and survey by Minoji and Sakai (2024).

Raishan has been far more neglected in terms of academic study, commercialisation, and policy initiatives than the African fonio millets. As part of this study, authors were able to contact and collaborate with researchers at the Botanical Study of India, and with Dr. Meera Das and Fullmerries Puwein at the North-Eastern Hill University, in the Shillong region, Meghalaya, India. Communication with rural farmers in the Khasi Hills has confirmed that the crop was still being cultivated in early 2024, though it is rare and under threat, and being replaced by varieties of higher-yielding finger millet (*Eleusine coracana* (L.) Gaertn.). Raishan has a host of traits that may be beneficial and interesting for further study, including its large grains and long, numerous racemes - and may provide useful information to breeding programmes to benefit food security in both West Africa and India, if international effort is made to protect this species. Recent Indian government initiatives including the UN’s International Year of Millets 2023 made numerous references to raishan in advertising material and policy documents, but there has so far been little effort to accelerate research or develop initiatives for its conservation and promotion. Seed-banking would be a particularly effective method for its protection.

#### Clade 6 – Polish millet, *D. ciliaris*, and relatives

The most noticeable difference in topology between trees is the placement of *D. ciliaris* and relatedness to *D. sanguinalis* and *D. compacta*. Henrard (1950) makes observations about the blurred and overlapping suite of characters these three species display, and their overlapping occurrences throughout Asia, Europe, and northern Africa. Genetic study of these species has supported two discrete, separate species in *D. sanguinalis* and *D. ciliaris*, but with closely shared ancestry (Adoukonou-Sagbadja *et al*., 2010).

*Digitaria sanguinalis* is estimated as both triploid and tetraploid in this study, corroborating other literature (Šmarda et al., 2014), and also reported as hexaploid and above elsewhere (2n=54, Xue et al., 1992; Bennet et al., 1998). *D. ciliaris* is similarly estimated to be triploid and tetraploid here, and as tetraploid and hexaploid in literature (Carnahan and Hill, 1961; Trouin, 1972).

It may be interpreted from the incongruence between the ML and ASTRAL-III trees, and from their complex genomic history (with polyploids above hexaploidy having been observed), that there is likely to be introgression or reticulation events between the two species and close relatives, as morphological traits vary and mix across the distribution of both species. The closely related species *D. sanguinalis*, *D. ciliaris*, *D. nuda*, and *D. horizontalis*, are known to frequently occupy fields together (Da Silva *et al*. 2024). It may be useful to consider clade 6 containing these species in the ASTRAL-III tree as a broad taxonomic complex. Somewhere in this complex, a landrace precursor in populations selected as Polish millet would have been foraged and cultivated in Europe – likely a hybrid between several of the species in this clade. This contrasts the well-resolved and distinct lineages of the other *Digitaria* crops. As *D. sanguinalis* is no longer cultivated as a food crop in Europe, it is difficult to untangle this complex - the complicated polyploid nature of this clade also makes it challenging to conduct genomic studies. These issues make it difficult to predict how useful reviving and utilising the cultivation of Polish millet may be to sustainable agricultural programmes in the future.

### Polyploidy

While the binate clade in this study does not contain significantly more polyploid species than the ternate clade, the species *D. sanguinalis* and *D. ciliaris* in the binate clades are the only *Digitaria* which are commonly reported to be hexaploid or above. Inconsistencies between chromosome count studies in the literature and some of the ploidy estimations in this study (especially around *D. exilis* wild relatives and *D. radicosa*) are concerning, and show how challenging it can be to conduct phylogenetic studies of polyploid species using NGS target-capture sequencing data (Kyriakidou *et al*., 2018; Dufresne et al., 2013; Griffin et al., 2011). There is certainly scope for future research to reassess the complex interactions of *Digitaria* species topologies, crop domestication, and polyploidy in Panicoid grasses, using methods such as that of Masters *et al*. (2024) for *Brachiaria* grasses using allele phasing. Phylogenetic network analysis may also provide a useful method to detect possible reticulation and hybridisation events, which would be valuable to apply to *Digitaria* grasses (Solís-Lemus and Ané, 2016; Solís-Lemus et al., 2017).

### Evolution and Biogeography

From species distributions and BEAST node age estimation it is predicted that early groups of *Digitaria* diverged at least 15-17million years ago in central or South-Eastern Africa, during the Miocene period. This corroborates with estimated divergences of Anthephorinae (the subtribe within which *Digitaria* belongs) dated to a mean age of 18.43mya by Gallaher *et al*. (2022), with a likelihood of ∼21% having originated in the Afrotropics region. The tribe Paniceae are estimated to 29.96mya with a 38% likelihood of origin in the Afrotropics, in the same study. Hackel *et al*. (2018) estimates the divergence of Paniceae at around 22mya, and the split between binate and ternate *Digitaria* at around 15mya (similar to 15-17mya in this study). Africa has generally been suggested as an area of early divergences for many grass lineages (Gallaher *et al*., 2022; Peterson and Soreng, 2022), especially C4 grasses, as well as for Poaceae as a whole (Bouchenak-Khelladi *et al*., 2010), which corroborates our results here.

In the phylogenetic trees, the PCA, and biogeographic and time analysis, it appears that *D. monodactyla*, *D. debilis*, *D. leptorhachis*, *D. humbertii*, and *D. compressa* are the earliest diverging and most genetically dissimilar to other later diverging *Digitaria,* and are all African native species. *D. monodactyla* is one of the only binate species within the ternate group, and has uncommonly single, spicate racemes. *D. monodactyla* and *D. compressa* also have fire-adaptive culm crowns, adapted to fire-regime lifestyles in seasonal savannah open grasslands in south-eastern Africa (see photos of growth forms, Figure S7). The mid-late Miocene saw aridification of parts of south-eastern Africa which shaped the evolution of plant species (Osborne, 2008; Pokorny *et al*., 2015), and in the later period allowed for the expansion of savannah and grasslands (Sepulchre *et al*., 2006). The formation of these grasslands saw the dominance of C4 grassland species in these drier climates between 20-10mya (Peppe *et al*., 2023; Edwards *et al*., 2010), which are thought to have coincided with ‘enhanced fire activity’ (Hoetzel *et al*., 2013; Karp *et al*., 2018). This expansion of C4 grasses, increasingly arid climate, and fire activity would have both shaped and supported the diversification of *Digitaria* grasses possibly including ancestors of the early diverging extant species *D. monodactyla* and *D. compressa*, and the spread of both the ternate fonio lineage to spread West, and the binate raishan and Polish millet lineages to spread north to Europe and Asia.

In other phylogenetic studies including *Digitaria*, the genera *Chaetopoa* C.E.Hubb., *Chlorocalymma* Clayton, *Anthephora* Schreb, and *Baptorhachis* Clayton & Renvoize are embedded exclusively within the binate clade, and not among the ternate species clades (Grass Working Group III, 2024; Hackel *et al*., 2018; Touafchia *et al*., 2023), including: *Chaetopoa taylorii* C.E.Hubb, *Chlorocalymma cryptacanthum* Clayton, *Taeniorhachis repens* Cope, *Anthephora pubescens* Nees, *A. cristata* (Döll) Hack. ex De Wild. & T.Durand, *Tarigidia aequiglumis* (Gooss.) Stent, *Stereochlaena caespitosa* Clayton, and *Baptorhachis foliacea* (Clayton) Clayton. These species are also native to dryland African grasslands (POWO, https://powo.science.kew.org/). *Chaetopoa taylorii*, *Chlorocalymma cryptacanthum*, *Baptorhachis foliacea*, and *Anthephora pubescens* also all have single racemes (similar to the spicate inflorescence of *Digitaria monodactyla*). These species provide an ‘African grassland’ core to the otherwise quite widely distributed binate species – and possibly represent a closer relationship to the earlier diverging *Digitaria* in clades 1 and 4 from south-eastern Africa, with close morphological similarity and ecological habitat.

While the ternate clades appear to be more morphologically uniform, and have undergone more simple dispersals mainly within Africa, there appears to be more complicated movement within the binate group, especially around the Arabian peninsula, likely between Madagascar and India, and between Asia and Europe. These movements give rise to inconsistencies in both phylogenetic and morphological systematics, and challenges to genomic assembly posed by polyploidy. These species cause the most taxonomic problems, as described above with *D. sanguinalis* and *D. ciliaris*. Genetic reticulations are common across the Paniceae (Grass Working Group III 2024), and recent genomic work by Minoji and Sakai (2024) comments on the difficulty of assembling references for polyploid *Digitaria* species.

From the few native species sequenced as part of this study, *Digitaria* in Madagascar appear to have occured between 10-15mya, with the separation of *D. humbertii* (endemic to Madagascar) from *D. debilis* and *D. leptorhachis* (occurring in mainland Africa, and *D. debilis* also occurring in Madagascar). *D. humbertii* is a grazing-adapted grassland species, with rhizomes, spreading and mat-forming – its sister species *D. debilis* and *D. leptorhachis* occupy a similar ecological niche, but are more erect in their growth habit. In a study by Randrianarimanana *et al*. (2024), ethnobotanic interviews with farmers across three regions in central and eastern Madagascar, *D. humbertii* was cited as the most troublesome grass weed in agricultural settings, followed by *D. longiflora*, which also has a similar growth habit to *D. humbertii*. Other *Digitaria* including *D. ternata*, *D. ciliaris* and *D. sanguinalis*, and *D. debilis* were also reported. This ‘weedy’ habit is seen in early and late diverging species on mainland Africa and Madagascar, and this phenotype was shown by Touafchia *et al*. (2023) to not be specific to any clade across the genus, and to have instead evolved convergently many times. Hackel *et al*. (2018) in investigating the origin and diversity of C3 and C4 grass evolutions in Madagascar, comments that endemic C4 Panicoid grasses are likely to have appeared during the Miocene – supporting the speciation of the *D. humbertii*-*D. debilis* clade during this period.

### Fonio Domestication

Despite a long history of evolution, *Digitaria* crops do not show significant developments that make them overly distinct from their wild relatives: there is no distinct abscission zone change to suggest a strongly evolved non-shattering phenotype, the grain sizes overlap, and there is only complete loss of trichomes in *D. exilis* (the most commonly cultivated species). In a study of 203 global food crops by Meyer *et al*. (2012), the term ‘semi-domesticated’ is used to describe crops which have achieved only partial differentiation from wild species, and seems relevant to apply to *Digitaria* crops here.

The only study to consider fonio abscission zones (a key mechanism for regulating shattering) in depth is Patterson *et al*. (2016), where an SEM figure of the *Digitaria exilis* abscission zone is shown, but without comparison to wild *Digitaria*. It shows that the abscission zone for *Digitaria* can be found at the base of the spikelet, similar to other closely related millet crops. This can be seen in our results, with both the shattered and complete spikelets. Abscission zones are known to form in differently ways across the grasses, with differential cell size and lignification being two strategies for formation, regulated by genes including *sh1*, *sh4*, *qSH3*, and *qShi1* (Mascher *et al*., 2024; Yu and Kellogg, 2018; Yu *et al*., 2020). In sorghum, there have been three different mutations of the *sh1* gene on chromosome 1 (Fuller and Stevens, 2018), which deactivate the formation of the abscission zone, and reduce shattering. Differential cell shapes appear to be important to abscission zone formation in *Digitaria*, though there is no obvious difference in formation between wild and cultivated species. In the assembly of the fonio genome (with surveys for domestication signatures), Abrouk *et al*. (2020) find that only 37% of accessions of fonio across its whole cultivated range have a deletion in the *sh1* gene, and that plants with the deletion only show a 7% chance of reduced shattering. Although this deletion has not been reported at all in the wild relative *D. longiflora*, this is a low expression of non-shattering phenotypes even with the gene under selection.

In a survey of major plant domestication pathways (Fuller *et al*., 2023), the domestication centre of white fonio is placed in the Inner Niger Basin, Mali, with African rice, pearl millet, and cowpea, but only fonio is described as having been domesticated through a ‘segetal pathway’, versus a ‘grain pathway’. Segetal is defined as “former weedy species that grew in agricultural contexts that were added to the crop repertoire”. These segetal domesticates, which grew as weeds and were adopted into mainstream cultivation through convenience, were thought to be domesticated much later, and more quickly than others, likely around 2000-3000 years ago. Other ‘segetal’ pathway crops including rye and oats probably grew as weeds in the fields of wheat and barley before selection. For fonio, a suggestion is that these harbour crops would have been African rice and pearl millet in the Mali region. Pearl millet was domesticated much earlier in Mali, and was very popular until African rice, fonio and sorghum were later incorporated into local food selection (Champion *et al*., 2021). This explains why fonio may have poorly developed domestication traits. A genetic resources and breeding programme study of fonio in Nigeria and Benin (Kanlindogbe *et al*., 2020) agrees that across West Africa there has been little conscious, targeted selection of fonio cultivars, describing cultivated varieties as ‘ecotypes’. In Burton et al. (2024), fonio farmers interviewed in Guinea explained that landraces were only selected for suitability to local microclimates which changed in rotations every few years, and not for specific uses, suggesting a low domestication pressure for morphological traits, and explaining why traits like shattering have not been significantly improved. Another factor may be the co-existence of *Digitaria exilis* and *D. longiflora* often in the same fields, noted by Burton et al., (2024) in Guinea, and with overlapping occurrence ranges across West Africa (POWO, 2025). Though low (around 2%; Barnaud et al., 2017), outcrossing in *D. exilis* plants has been observed, and introgressions between crop and wild species are still likely, effectively slowing the rate of domestication pressure. This adds weight to the theory that cereal domestication can be a long, slow, ongoing landscape-level process, with no single epicenter for some species (Allaby et al., 2017; Allaby et al., 2022).

## Conclusions

This study presents a comprehensive, widely-sampled and well-resolved phylogeny of the grass genus *Digitaria*, including all 4 lineages domesticated for food crops (and their wild relatives) throughout history and geographic regions. Tree topologies are largely congruent with classification and grouping of species by previous studies, and 3 domesticated crops (white and black fonio, and raishan) appear as distinct, monophyletic lineages, while the European Polish millet appears to be part of a complicated genetic and taxonomic complex with *D. ciliaris* and *D. velutina*. Our results support that four *Digitaria* crops were produced through separate evolutionary lineages. Type specimens of the rare crops Indian endemic raishan and West African black fonio, are sequenced and presented here in phylogenetic analyses for the first time. The closest wild relatives to raishan are found to be wild Asian native species *D. cruciata*, *D. radicosa* and *D. setigera*. New close relatives for both fonio species are identified in *D. atrofusca* (for black fonio), and *D. fuscescens* and *D. argyrotricha* (for white fonio), all African native species. We also confirm through communication with specialists in India and Nigeria that *D. iburua* and *D. compacta* were still cultivated in 2024.

Biogeographic analyses suggest an early origin of *Digitaria* in arid grasslands of South and Eastern Africa, with later dispersals and further speciation in West Africa, and to Asia and Europe over the course of the past 15 million years. Early extant *Digitaria* lineages include African species *D. monodactyla*, *D. compressa*, *D. gayana*, and *D. debilis*, *D. leptorhachis*, and *D. humbertii*. Both separate fonio lineages are estimated to have diverged from relatives between 2-3mya, raishan more recently, and *D. sanguinalis* also between 6 and 2mya. Fonio species were only seen to have experienced semi-domestication, experiencing loss of trichomes and developing longer racemes, but not significantly decreased shattering or abscission zone differentiation.

As the degradation of African grasslands - including large areas of millet cultivation - accelerates due to human activity and climate change, causing erratic weather patterns and increased fire activity, it will be vital to invest in crops that are resilient to fire and extreme climates (Osborne *et al*., 2018; Joseph *et al*. 2024; Knowles *et al*. 2025). *Digitaria* species, not only white and black fonio, have the evolutionary history and genetics likely to hold potential for improving food security for vulnerable areas in the future. Collections of fresh and dried plant and seed material for both raishan and black fonio alongside ethnobotanic data should also be prioritised and conserved, to prevent the loss of these rare traditional crops.

## Acknowledgements

This study is a part of G. Burton’s wider PhD project on fonio and food security, and we are thankful for funding from the United Kingdom Research and Innovation (UKRI) and the Natural Environment Research Council (NERC), administered through the Science and Solutions for a Changing Planet DTP, with help provided by Christiane Morgan at the Grantham Institute (Imperial College London), and Felix Forest at RGB Kew. Thank you to the NERC Environmental Omics Facility (NEOF) and Jemima Brinton for facilitating the sequencing of a *Digitaria sanguinalis* genome, which was useful to this study, and to the Leibniz-Institut Genebank (IPK) for providing germplasm. The authors would like to acknowledge and thank collaborators and staff at RBG Kew including curator-botanist Martin Xanthos for assisting with sampling the herbarium collection for sequencing, Sidonie Bellot, Alex Zuntini, George Tiley, Toby Kellogg, Mark Nesbitt, and Rafael Felipe de Almeida for useful discussions about grass evolution and phylogenetic analysis, the Jodrell Laboratory team Laszlo Csiba, Robyn S. Cowan, and Dion Devey, for helping to extra DNA from samples, Jan Hackel for discussions and help with lab work, and Lucy T. Smith for illustrations. Thank you also to the ADCRA team at Kew including Rafal Gutaker, Caspar Chater, and Amy Jackson for your help and advice on data analysis, and ongoing support to the lead author. We would like to thank researchers Nantenaina Rakotomalala and Fenitra Randrianarimanana at KMCC, Madagascar for assistance working with Malagasy *Digitaria*. In India, we would like to thank associate professor Meera Das and student Fullmerries Puwein at the North-Eastern Hill University, Shillong, and Harmindir Singh at the Botanical Survey of India, for correspondence about the crop raishan. Thanks to Abubakar Bello and students Ishaq Muawiyya, Ibrahim Tahir, at the Umaru Musa Yaradua University, Katsina, Nigeria, for your collaboration researching black fonio. Thank you also to Ryohei Terauchi and Toshiyuki Sakai at the Crop Evolution Laboratory, Kyoto, Japan, for your discussions on *Digitaria* evolution and breeding.

## Supplementary Data

Figure S1 – Morphological features of *Digitaria* grasses. From top left clockwise: inflorescence and racemes; spikelets; rachis and pedicels; and growth forms.

Figure S2 – Photo of a farmer in the Khasi Hills, India, with the millet raishan (*Digitaria compacta*). Reproduced from Singh & Arora (1972).

Figure S3 – Morphological traits including spikelet arrangements (binate vs. ternate) in a. and spikelet hair type (complex, simple or mostly glabrous) in b., across ML phylogeny of *Digitaria* species.

Figure S4 – Ploidy estimations produced by nQuire, given as 2 (diploid), 3 (triploid) and 4 (tetraploid), across ML phylogeny.

Figure S5 – Ancestral biogeographic origin of *Digitaria* species, produced by BioGeoBEARS analysis. Lineages are coloured by regions of native distribution from A-E. Clades corresponding to the four crop species are indicated.

Figure S6 - Scanning Electron Microscope photos of *Digitaria* abscission zones, wild species. Captured at Kew Gardens and the Natural History Museum (UK). Red arrows indicate abscission zone. From top left to bottom right: *D. abyssinica* (Schimper 82), *D. argyrotricha* (Scholz, H. 1096), *D. compressa* (A. Blair Rains 207), *D. debilis* (X.M. van der Burgt 1771)*, D. diagonalis* (I. Darbyshire 471), *D. gayana* (F.F. Merklinger 15), *D. perrotettii* (F.F. Merklinger 104), and *D. setigera* (Clayton 5338).

Figure S7 – Growth form morphologies of *Digitaria* species, for cultivated crops (row 1), wild relatives (rows 2 and 3), and early diverging fire-adapted species (row 4). Photos are to same scale, using scale bar in bottom left image.

Table S1 – Details of specimens sampled for DNA extraction and phylogenetic analysis. Table S2 – DNA sequence recovery statistics from HybPiper assembly analysis.

## Literature Cited

Abdoul-Raouf SM, Ju Q, Jianyu M, Zhizhai L. 2017. Utilization of wild relatives for maize (Zea mays L.) improvement. African Journal of Plant Science 11: 105–113.

Abrouk M, Ahmed HI, Cubry P, Šimoníková D, Cauet S, Pailles Y, Bettgenhaeuser J, Gapa L, Scarcelli N, Couderc M, et al. 2020. Fonio millet genome unlocks African orphan crop diversity for agriculture in a changing climate. Nature Communications 11: 4488.

Adoukonou-Sagbadja H, Wagner C, Dansi A, Ahlemeyer J, Daïnou O, Akpagana K, Ordon F, Friedt W. 2007. Genetic diversity and population differentiation of traditional fonio millet (Digitaria spp.) landraces from different agro-ecological zones of West Africa. Theoretical and Applied Genetics 115: 917–931.

Adoukonou-Sagbadja H, Wagner C, Ordon F, Friedt W. 2010. Reproductive System and Molecular Phylogenetic Relationships of Fonio Millets (Digitaria spp., Poaceae) with Some Polyploid Wild Relatives. Tropical Plant Biology 3: 240–251.

Allaby RG, Stevens CJ, Kistler L, Fuller DQ. 2022. Emerging evidence of plant domestication as a landscape-level process. Trends in Ecology & Evolution 37: 268–279.

Allaby RG, Stevens C, Lucas L, Maeda O, Fuller DQ. 2017. Geographic mosaics and changing rates of cereal domestication. Philosophical Transactions of the Royal Society B: Biological Sciences 372: 20160429.

Animasaun DA, Adedibu PA, Olawepo GK, Oyedeji S. 2023. Fonio millets. In: Neglected and Underutilized Crops. Elsevier, 201–219.

Baker WJ, Dodsworth S, Forest F, Graham SW, Johnson MG, McDonnell A, Pokorny L, Tate JA, Wicke S, Wickett NJ. 2021. Exploring Angiosperms353: An open, community toolkit for collaborative phylogenomic research on flowering plants. American Journal of Botany 108: 1059–1065.

Bennett M. 1998. DNA Amounts in Two Samples of Angiosperm Weeds. Annals of Botany 82: 121–134.

Bolger AM, Lohse M, Usadel B. 2014. Trimmomatic: a flexible trimmer for Illumina sequence data. Bioinformatics 30: 2114–2120.

Boonsuk B, Chantaranothai P, Hodkinson TR. 2016. A taxonomic revision of the genus Digitaria (Panicoideae: Poaceae) in mainland Southeast Asia. Phytotaxa 246: 248.

Borowiec ML. 2016. AMAS: a fast tool for alignment manipulation and computing of summary statistics. PeerJ 4: e1660.

Bouchenak-Khelladi Y, Verboom GA, Savolainen V, Hodkinson TR. 2010. Biogeography of the grasses (Poaceae): a phylogenetic approach to reveal evolutionary history in geographical space and geological time: ANCESTRAL BIOGEOGRAPHY AND ECOLOGY OF GRASSES. Botanical Journal of the Linnean Society 162: 543–557.

Bouckaert R, Heled J, Kühnert D, Vaughan T, Wu C-H, Xie D, Suchard MA, Rambaut A, Drummond AJ. 2014.BEAST 2: A Software Platform for Bayesian Evolutionary Analysis (A Prlic, Ed.). PLoS Computational Biology 10: e1003537.

Brush S, Kesseli R, Ortega R, Cisneros P, Zimmerer K, Quiros C. 1995. Potato Diversity in the Andean Center of Crop Domestication. Conservation Biology 9: 1189–1198.

Burton G, Gori B, Camara S, Ceci P, Conde N, Couch C, Magassouba S, Vorontsova MS, Ulian T, Ryan P. 2024. Landrace diversity and heritage of the indigenous millet crop fonio ( *Digitaria exilis* ): Socio-cultural and climatic drivers of change in the Fouta Djallon region of Guinea. PLANTS, PEOPLE, PLANET: ppp3.10490.

Camacho C, Coulouris G, Avagyan V, Ma N, Papadopoulos J, Bealer K, Madden TL. 2009. BLAST+: architecture and applications. BMC Bioinformatics 10: 421.

Carnahan HL, Hill HD. 1961. Cytology and genetics of forage grasses. The Botanical Review 27: 1–162.

Champion L, Fuller DQ, Ozainne S, Huysecom É, Mayor A. 2021. Agricultural diversification in West Africa: an archaeobotanical study of the site of Sadia (Dogon Country, Mali). Archaeological and Anthropological Sciences 13: 60.

Charif D, Lobry JR. 2007. SeqinR 1.0-2: A Contributed Package to the R Project for Statistical Computing Devoted to Biological Sequences Retrieval and Analysis. In: Bastolla U, Porto M, Roman HE, Vendruscolo M, eds. Biological and Medical Physics, Biomedical Engineering. Structural Approaches to Sequence Evolution. Berlin, Heidelberg: Springer Berlin Heidelberg, 207–232.

Charmet G. 2011. Wheat domestication: Lessons for the future. Comptes Rendus. Biologies 334: 212–220.

Clayton. 2016. Clayton W, Vorontsova M, Harman T, Williamson H. 2016. GrassBase - The Online World Grass Flora. GrassBase-The Online World Grass Flora.

Cruz J-F, Béavogui F, Dramé D, Diallo TA. 2016. Fonio, an African cereal. Montpellier] [Conakry (Guinée): Cirad [UMR Qualisud] IRAG, Institut de recherche agronomique de Guinée.

Dempewolf H, Baute G, Anderson J, Kilian B, Smith C, Guarino L. 2017. Past and Future Use of Wild Relatives in Crop Breeding. Crop Science 57: 1070–1082.

Doyle & Doyle. 1987. Doyle, J.J., Doyle, J.L., “A rapid DNA isolation procedure for small quantities of fresh leaf tissue,” Phytochemical Bulletin, 19 (1). 11–15. 1987.

Dufresne F, Stift M, Vergilino R, Mable BK. 2014. Recent progress and challenges in population genetics of polyploid organisms: an overview of current state-of-the-art molecular and statistical tools. Molecular Ecology 23: 40–69.

Edwards EJ, Osborne CP, Strömberg CAE, Smith SA, C Grasses Consortium, Bond WJ, Christin P-A, Cousins AB, Duvall MR, Fox DL, et al. 2010. The Origins of C_4_ Grasslands: Integrating Evolutionary and Ecosystem Science. Science 328: 587– 591.

Fatima Z, Ahmed M, Hussain M, Abbas G, Ul-Allah S, Ahmad S, Ahmed N, Ali MA, Sarwar G, Haque EU, et al. 2020. The fingerprints of climate warming on cereal crops phenology and adaptation options. Scientific Reports 10: 18013.

Fuller DQ, Denham T, Allaby R. 2023. Plant domestication and agricultural ecologies. Current Biology 33: R636–R649.

Fuller DQ, Stevens CJ. 2018.Sorghum Domestication and Diversification: A Current Archaeobotanical Perspective | SpringerLink.

Gallaher TJ, Peterson PM, Soreng RJ, Zuloaga FO, Li D, Clark LG, Tyrrell CD, Welker CAD, Kellogg EA, Teisher JK. 2022. Grasses through space and time: An overview of the biogeographical and macroevolutionary history of Poaceae. Journal of Systematics and Evolution 60: 522–569.

Gilani, Syed & Khan, Mir & Shinwari, Zabta & Hussain, Farrukh & Yousaf, Zubaida. 2003. Taxonomic Relationship of the Genus Digitaria in Pakistan. Pakistan Journal of Botany. 35. 261–278. ResearchGate.

Glémin S, Bataillon T. 2009. A comparative view of the evolution of grasses under domestication. New Phytologist 183: 273–290.

Gould FW, Soderstrom TR. 1967. CHROMOSOME NUMBERS OF TROPICAL AMERICAN GRASSES. American Journal of Botany 54: 676–683.

Gould FW, Soderstrom TR. 1974. Chromosome numbers of some Ceylon grasses. Canadian Journal of Botany 52: 1075–1090.

Grass Phylogeny Working Group III. 2024. A nuclear phylogenomic tree of grasses (Poaceae) recovers current classification despite gene tree incongruence. New Phytologist: nph.20263.

Griffin PC, Robin C, Hoffmann AA. 2011. A next-generation sequencing method for overcoming the multiple gene copy problem in polyploid phylogenetics, applied to Poa grasses. BMC Biology 9: 19.

Hackel J, Vorontsova MS, Nanjarisoa OP, Hall RC, Razanatsoa J, Malakasi P, Besnard G. 2018. Grass diversification in Madagascar: In situ radiation of two large C_3_ shade clades and support for a Miocene to Pliocene origin of C_4_ grassy biomes. Journal of Biogeography 45: 750–761.

Hajjar R, Hodgkin T. 2007. The use of wild relatives in crop improvement: a survey of developments over the last 20 years. Euphytica 156: 1–13.

Haller. 1768. Haller. 1768, Hist. Stirp. Helv. 2: 244.

Harlan JR, De Wet JMJ, Price EG. 1973. Comparative Evolution of Cereals. Evolution 27: 311.

Henrard JT. 1950. Monograph of the Genus Digitaria: By J. T. Henrard. Leiden, Netherlands: Universitaire Pers Leiden. 999 pages (illus.). 1950. Agronomy Journal 42: 631–631.

Hitu KW, M’Ribu K, Liang H, Mandelbaum C. 1997. Fonio Millets: Ethnobotany, Genetic Diversity and Evolution. South African Journal of Botany 63: 185–190.

Hoetzel S, Dupont L, Schefuß E, Rommerskirchen F, Wefer G. 2013. The role of fire in Miocene to Pliocene C4 grassland and ecosystem evolution. Nature Geoscience 6: 1027–1030.

Hooker, J.D. 1897. Flora of British India. Vol 7, p.14.

Hoshino T, Davidse G. 1988. Chromosome Numbers of Grasses (Poaceae) From Southern Africa. I. Annals of the Missouri Botanical Garden 75: 866–873.

Johnson MG, Gardner EM, Liu Y, Medina R, Goffinet B, Shaw AJ, Zerega NJC, Wickett NJ. 2016. HybPiper: Extracting coding sequence and introns for phylogenetics from high-throughput sequencing reads using target enrichment. Applications in Plant Sciences 4: 1600016.

Johnson MG, Pokorny L, Dodsworth S, Botigué LR, Cowan RS, Devault A, Eiserhardt WL, Epitawalage N, Forest F, Kim JT, et al. 2019. A Universal Probe Set for Targeted Sequencing of 353 Nuclear Genes from Any Flowering Plant Designed Using k-Medoids Clustering (S Renner, Ed.). Systematic Biology 68: 594–606.

Jones EAL, Contreras DJ, Everman WJ. 2021. Digitaria ciliaris, Digitaria ischaemum, and Digitaria sanguinalis. In: Biology and Management of Problematic Crop Weed Species. Elsevier, 173–195.

Joseph GS, Seymour CL, Rakotoarivelo AR. 2024. Fire incongruities can explain widespread landscape degradation in Madagascar’s forests and grasslands. *PLANTS, PEOPLE*, PLANET 6: 656–669.

Junier T, Zdobnov EM. 2010. The Newick utilities: high-throughput phylogenetic tree processing in the U nix shell. Bioinformatics 26: 1669–1670.

Kanlindogbe C, Sekloka E, Kwon-Ndung EH. 2020. Genetic Resources and Varietal Environment of Grown Fonio Millets in West Africa: Challenges and Perspectives. Plant Breeding and Biotechnology 8: 77–88.

Karp AT, Behrensmeyer AK, Freeman KH. 2018. Grassland fire ecology has roots in the late Miocene. Proceedings of the National Academy of Sciences 115: 12130– 12135.

Kashyap A, Garg P, Tanwar K, Sharma J, Gupta NC, Ha PTT, Bhattacharya RC, Mason AS, Rao M. 2022. Strategies for utilization of crop wild relatives in plant breeding programs. Theoretical and Applied Genetics 135: 4151–4167.

Katoh K, Standley DM. 2013. MAFFT Multiple Sequence Alignment Software Version 7: Improvements in Performance and Usability. Molecular Biology and Evolution 30: 772–780.

Kellogg EA. 2015. Inflorescence Structure. In: Flowering Plants. Monocots. Cham: Springer International Publishing, 25–38.

Knowles T, Stevens N, Amoako EE, Armani M, Barbosa C, Beale C, Bond W, Chidumayo E, Courtney-Mustaphi C, Dintwe K, et al. 2025. Viability and desirability of financing conservation in Africa through fire management. Nature Sustainability.

Koul KK, Gohil RN. 1991. Cytogenetic Studies on Some Kashmir Grasses. VIII. Tribe Agrostideae, Festuceae and Paniceae.: Tribe Agrostideae, Festuceae and Paniceae. CYTOLOGIA 56: 437–452.

Kyriakidou M, Tai HH, Anglin NL, Ellis D, Strömvik MV. 2018. Current Strategies of Polyploid Plant Genome Sequence Assembly. Frontiers in Plant Science 9.

Lo Medico JM, Tosto DS, Rua GH, De Agrasar ZER, Scataglini MA, Vega AS. 2017. Phylogeny of *Digitaria* Sections *Trichachne* and *Trichophorae* (Poaceae, Panicoideae, Paniceae): A Morphological and Molecular Analysis. New Circumscription and Synopsis. Systematic Botany 42: 37–53.

Löve Á. 1985. Chromosome Number Reports LXXXVI. Taxon 34: 159–164.

Mabhaudhi T, Chimonyo VGP, Hlahla S, Massawe F, Mayes S, Nhamo L, Modi AT. 2019. Prospects of orphan crops in climate change. Planta 250: 695–708.

Mai U, Mirarab S. 2018. TreeShrink: fast and accurate detection of outlier long branches in collections of phylogenetic trees. BMC Genomics 19: 272.

Mascher M, Marone MP, Schreiber M, Stein N. 2024. Are cereal grasses a single genetic system? Nature Plants 10: 719–731.

Masters LE, Tomaszewska P, Schwarzacher T, Hackel J, Zuntini AR, Heslop-Harrison P, Vorontsova MS. 2024. Phylogenomic analysis reveals five independently evolved African forage grass clades in the genus *Urochloa*. Annals of Botany 133: 725–742.

Matzke NJ. 2013. Probabilistic historical biogeography: new models for founder-event speciation, imperfect detection, and fossils allow improved accuracy and model-testing. Frontiers of Biogeography 5.

Mehra PN. 1982. Cytology of East Indian grasses. Chandigarh, India: P. N. Mehra.

Meyer RS, DuVal AE, Jensen HR. 2012. Patterns and processes in crop domestication: an historical review and quantitative analysis of 203 global food crops. New Phytologist 196: 29–48.

Minh BQ, Schmidt HA, Chernomor O, Schrempf D, Woodhams MD, Von Haeseler A, Lanfear R. 2020. IQ-TREE 2: New Models and Efficient Methods for Phylogenetic Inference in the Genomic Era (E Teeling, Ed.). Molecular Biology and Evolution 37: 1530–1534.

Minoji K, Sakai T. 2024.A chromosome-scale genome assembly of Timorese crabgrass ( *Digitaria radicosa* ): a useful genomic resource for the Poaceae (I Parkin, Ed.). G3: Genes, Genomes, Genetics: jkae242.

Morrone O, Aagesen L, Scataglini MA, Salariato DL, Denham SS, Chemisquy MA, Sede SM, Giussani LM, Kellogg EA, Zuloaga FO. 2012. Phylogeny of the Paniceae (Poaceae: Panicoideae): integrating plastid DNA sequences and morphology into a new classification. Cladistics 28: 333–356.

Netolitzky. 1914. Die Hirse aus antiken Funden.

Ngom A, Gueye MC, Gueye M, Billot C, Calatayud C, Diop BM, Kane NA, Piquet M, Vigouroux Y, Zekraoui L, et al. 2019. Cross-species amplification of microsatellite loci developed *Digitaria exilis* Stapf in related *Digitaria* species. Journal of Applied Biosciences 129: 12982.

Nyam D, Eh K-N, Ap W. 2017. Genetic Affinity and Breeding Potential of Phenologic Traits of Acha (fonio) in Nigeria.

Osborne CP. 2008. Atmosphere, ecology and evolution: what drove the Miocene expansion of C_4_ grasslands? Journal of Ecology 96: 35–45.

Osborne CP, Charles-Dominique T, Stevens N, Bond WJ, Midgley G, Lehmann CER. 2018. Human impacts in African savannas are mediated by plant functional traits. New Phytologist 220: 10–24.

Paradis E, Schliep K. 2019. ape 5.0: an environment for modern phylogenetics and evolutionary analyses in R (R Schwartz, Ed.). Bioinformatics 35: 526–528.

Parsons JJ. 1972. Spread of African Pasture Grasses to the American Tropics. Journal of Range Management 25: 12.

Patterson SE, Bolivar-Medina JL, Falbel TG, Hedtcke JL, Nevarez-McBride D, Maule AF, Zalapa JE. 2016. Are We on the Right Track: Can Our Understanding of Abscission in Model Systems Promote or Derail Making Improvements in Less Studied Crops? Frontiers in Plant Science 6.

Peppe DJ, Cote SM, Deino AL, Fox DL, Kingston JD, Kinyanjui RN, Lukens WE, MacLatchy LM, Novello A, Strömberg CAE, et al. 2023. Oldest evidence of abundant C_4_ grasses and habitat heterogeneity in eastern Africa. Science 380: 173–177.

Pessoa-Filho M, Martins AM, Ferreira ME. 2017. Molecular dating of phylogenetic divergence between Urochloa species based on complete chloroplast genomes. BMC Genomics 18: 516.

Peterson PM, Soreng RJ. 2022. The biogeography of grasses (Poaceae). Journal of Systematics and Evolution 60: 473–475.

Pokorny L, Riina R, Mairal M, Meseguer AS, Culshaw V, Cendoya J, Serrano M, Carbajal R, Ortiz S, Heuertz M, et al. 2015. Living on the edge: timing of Rand Flora disjunctions congruent with ongoing aridification in Africa. Frontiers in Genetics 6.

Portères R. 1955. Les Céréales mineures du genre Digitaria en Afrique et en Europe (suite). Journal d’agriculture tropicale et de botanique appliquée 2: 477–510.

Portères R. 1957. Céréales mineures du genre Digitaria dans l’Inde. Journal d’agriculture tropicale et de botanique appliquée 4: 106–107.

Portères R. 1976. African Cereals: Eleusine, Fonio, Black Fonio, Tejf, Brachiaria, paspalum, Pennisetum, and African Rice. In: Harlan JR, Wet JMJD, Stemler ABL, eds. Origins of African Plant Domestication. DE GRUYTER MOUTON, 409–452.

POWO. 2025. Plants of the World Online. Kew Science. Plants of the World Online. https://powo.science.kew.org/

R Core Team. 2024. R Core Team (2024). _R: A Language and Environment for Statistical Computing_. R Foundation for Statistical Computing, Vienna, Austria. <https://www.R-project.org/>.

Rambaut A, Drummond AJ, Xie D, Baele G, Suchard MA. 2018. Posterior Summarization in Bayesian Phylogenetics Using Tracer 1.7 (E Susko, Ed.). Systematic Biology 67: 901–904.

Randrianarimanana NFH, Rakotomalala NH, MacKinnon L, Rakotoarinivo M, Randriamampianina J, Ralimanana H, Ryan P, Vorontsova MS. 2024. Local perceptions of the benefits versus negative impacts of weedy grasses in central Madagascar, with a focus on the genus *Digitaria*. *PLANTS, PEOPLE*, PLANET 6: 710–728.

Reeves. Save and Grow in practice: Maize, rice, wheat.

Revell LJ. 2012. phytools: an R package for phylogenetic comparative biology (and other things). Methods in Ecology and Evolution 3: 217–223.

Sepulchre P, Ramstein G, Fluteau F, Schuster M, Tiercelin J-J, Brunet M. 2006. Tectonic Uplift and Eastern Africa Aridification. Science 313: 1419–1423.

Singh HB, Arora RK. 1972. Raishan (Digitaria sp.)—a minor millet of the Khasi Hills, India. Economic Botany 26: 376–380.

Šmarda P, Bureš P, Horová L, Leitch IJ, Mucina L, Pacini E, Tichý L, Grulich V, Rotreklová O. 2014. Ecological and evolutionary significance of genomic GC content diversity in monocots. Proceedings of the National Academy of Sciences 111.

Smith SA, Brown JW, Walker JF. 2018.So many genes, so little time: A practical approach to divergence-time estimation in the genomic era (H Escriva, Ed.). PLOS ONE 13: e0197433.

Smith SA, Dunn CW. 2008. Phyutility: a phyloinformatics tool for trees, alignments and molecular data. Bioinformatics 24: 715–716.

Solís-Lemus C, Ané C. 2016. Inferring Phylogenetic Networks with Maximum Pseudolikelihood under Incomplete Lineage Sorting. PLOS Genetics 12: e1005896.

Solís-Lemus C, Bastide P, Ané C. 2017. PhyloNetworks: A Package for Phylogenetic Networks. Molecular Biology and Evolution 34: 3292–3298.

Stapf O. 1915. Iburu and Fundi, Two Cereals of Upper Guinea. (Digitaria iburua; D. exilis.). Bulletin of Miscellaneous Information (Royal Gardens, Kew) 1915: 381.

Stein JC, Yu Y, Copetti D, Zwickl DJ, Zhang L, Zhang C, Chougule K, Gao D, Iwata A, Goicoechea JL, et al. 2018. Genomes of 13 domesticated and wild rice relatives highlight genetic conservation, turnover and innovation across the genus Oryza. Nature Genetics 50: 285–296.

Thiers, B.M. 2025. Index Herbariorum. https://sweetgum.nybg.org/science/ih/.

Tikam K, Phatsara C, Mikled C, Vearasilp T, Phunphiphat W, Chobtang J, Cherdthong A, Südekum K-H. 2013. Pangola grass as forage for ruminant animals: a review. SpringerPlus 2: 604.

Touafchia S, Maurin O, Boonsuk B, Hodkinson TR, Chantaranothai P, Rakotomalala N, Randrianarimanana F, Randriamampianina JA, Roy S, MacKinnon L, et al. 2023. Evolutionary history, traits, and weediness in *Digitaria* (Poaceae: Panicoideae). Botanical Journal of the Linnean Society 203: 1–19.

Trouin M. 1972. Nombres chromosomiques de quelques Graminées du Soudan. Adansonia 12: 619–624.

Ulian T, Diazgranados M, Pironon S, Padulosi S, Liu U, Davies L, Howes MR, Borrell JS, Ondo I, Pérez-Escobar OA, et al. 2020. Unlocking plant resources to support food security and promote sustainable agriculture. *PLANTS, PEOPLE*, PLANET 2: 421–445.

Vega AS, Rua GH, Fabbri LT, Rúgolo De Agrasar ZE. 2009. A Morphology-Based Cladistic Analysis of *Digitaria* (Poaceae, Panicoideae, Paniceae). Systematic Botany 34: 312–323.

Virendra Kumar & B. Subramaniam. 1987. Chromosome atlas of flowering plants of the Indian subcontinent. Smithsonian Institution.

Wang L-G, Lam TT-Y, Xu S, Dai Z, Zhou L, Feng T, Guo P, Dunn CW, Jones BR, Bradley T, et al. 2020. Treeio: An R Package for Phylogenetic Tree Input and Output with Richly Annotated and Associated Data (S Kumar, Ed.). Molecular Biology and Evolution 37: 599–603.

Warburton ML, Crossa J, Franco J, Kazi M, Trethowan R, Rajaram S, Pfeiffer W, Zhang P, Dreisigacker S, Ginkel MV. 2006. Bringing wild relatives back into the family: recovering genetic diversity in CIMMYT improved wheat germplasm. Euphytica 149: 289–301.

Weiß CL, Pais M, Cano LM, Kamoun S, Burbano HA. 2018. nQuire: a statistical framework for ploidy estimation using next generation sequencing. BMC Bioinformatics 19: 122.

Wickham H. 2016. ggplot2. Cham: Springer International Publishing.

Woodhouse MR, Hufford MB. 2019. Parallelism and convergence in post-domestication adaptation in cereal grasses. Philosophical Transactions of the Royal Society B: Biological Sciences 374: 20180245.

Xue, B. S., Weng, R. F. and Zhang, M. Z. 1992. Chromosome numbers of Shanghai plants I.

Yu Y, Kellogg EA. 2018. Inflorescence Abscission Zones in Grasses: Diversity and Genetic Regulation. In: Roberts JA, ed. Annual Plant Reviews online. Wiley, 497– 532.

Yu Y, Leyva P, Tavares RL, Kellogg EA. 2020. The anatomy of abscission zones is diverse among grass species. American Journal of Botany 107: 549–561.

Yu G, Smith DK, Zhu H, Guan Y, Lam TT. 2017. ggtree : an r package for visualization and annotation of phylogenetic trees with their covariates and other associated data (G McInerny, Ed.). Methods in Ecology and Evolution 8: 28–36.

Zhang C, Rabiee M, Sayyari E, Mirarab S. 2018. ASTRAL-III: polynomial time species tree reconstruction from partially resolved gene trees. BMC Bioinformatics 19: 153.

Zheng X, Peng Y, Qiao J, Henry R, Qian Q. 2024. Wild rice: unlocking the future of rice breeding. Plant Biotechnology Journal 22: 3218–3226.

