## Supplementary Material for "Phylogenetics, evolution and biogeography of four *Digitaria* food crop lineages across West Africa, India, and Europe"

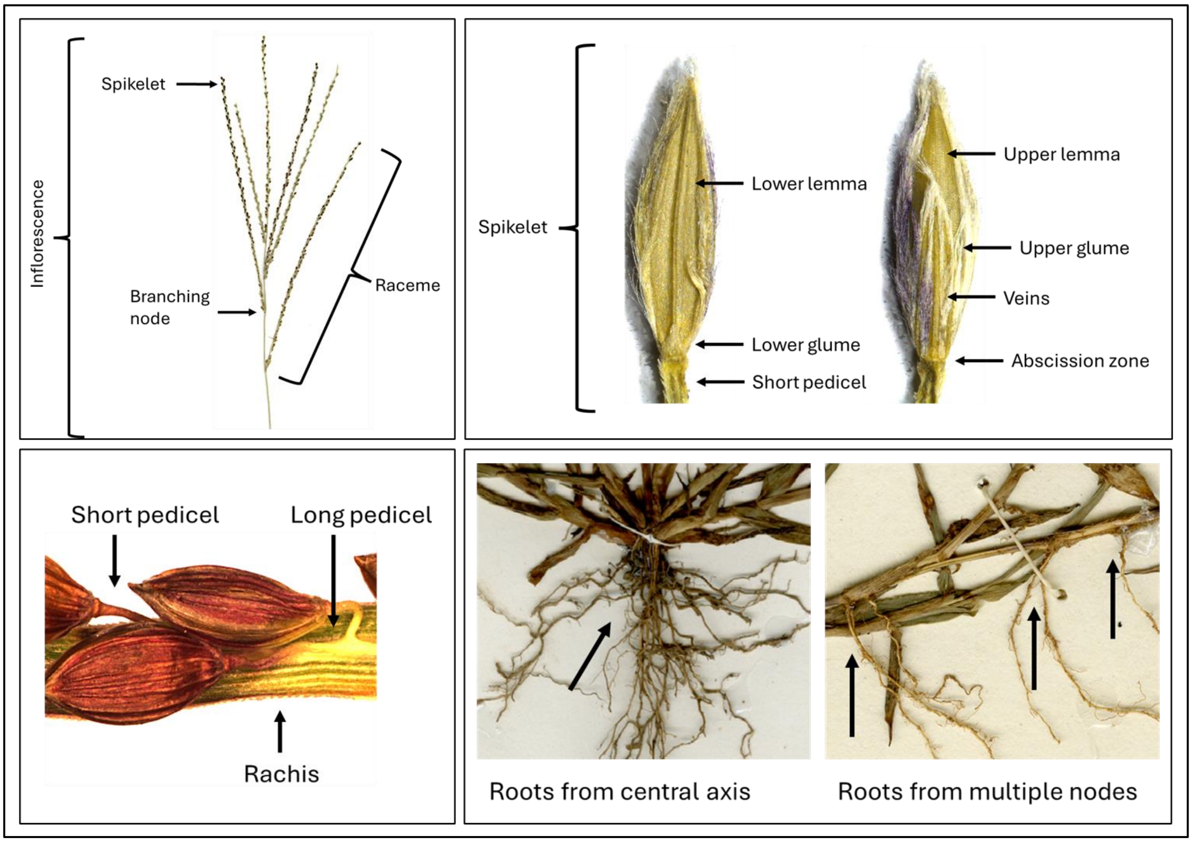


Figure S1 – Morphological features of *Digitaria* grasses. From top left clockwise: inflorescence and racemes; spikelets; rachis and pedicels; and growth forms.

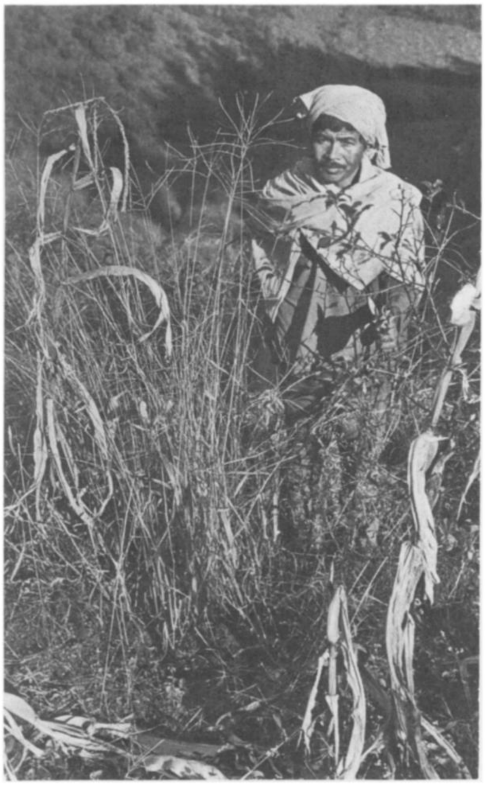

Figure S2 – Photo of a farmer in the Khasi Hills, India, with the millet raishan (*Digitaria compacta*). Reproduced from Singh & Arora (1972).

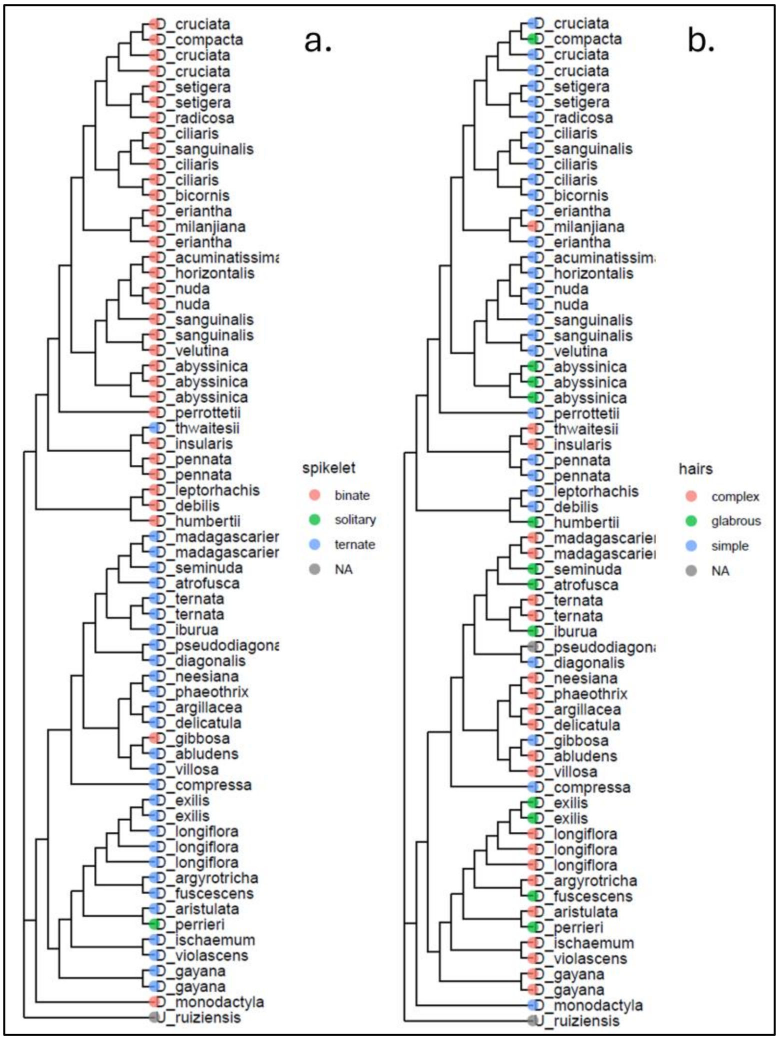


Figure S3 – Morphological traits including spikelet arrangements (binate vs. ternate) in a. and spikelet hair type (complex, simple or mostly glabrous) in b., across ML phylogeny of *Digitaria* species.

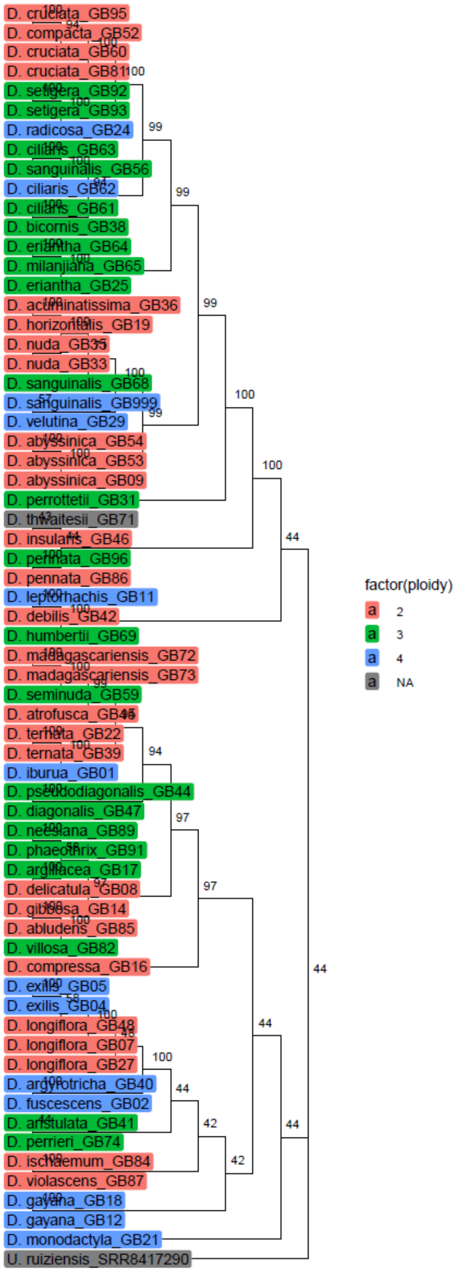


Figure S4 – Ploidy estimations produced by nQuire, given as 2 (diploid), 3 (triploid) and 4 (tetraploid), across ML phylogeny.

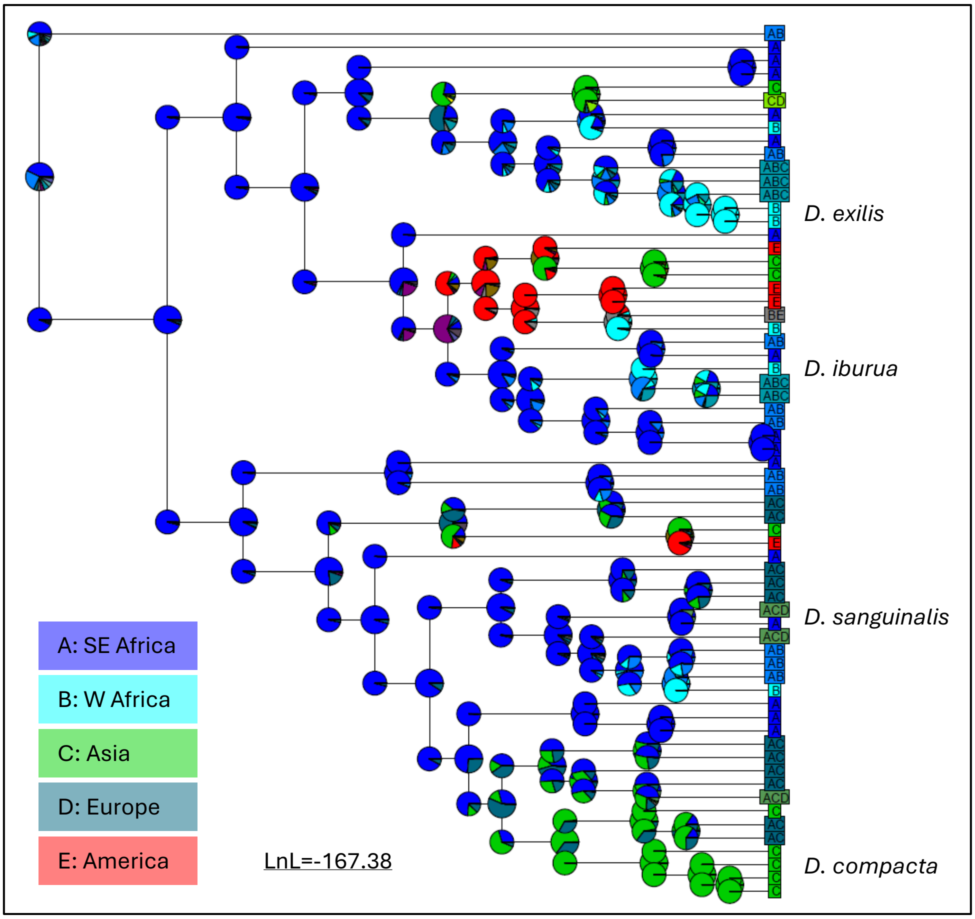


Figure S5 – Ancestral biogeographic origin of *Digitaria* species, produced by BioGeoBEARS analysis. Lineages are coloured by regions of native distribution from A-E. Clades corresponding to the four crop species are indicated.

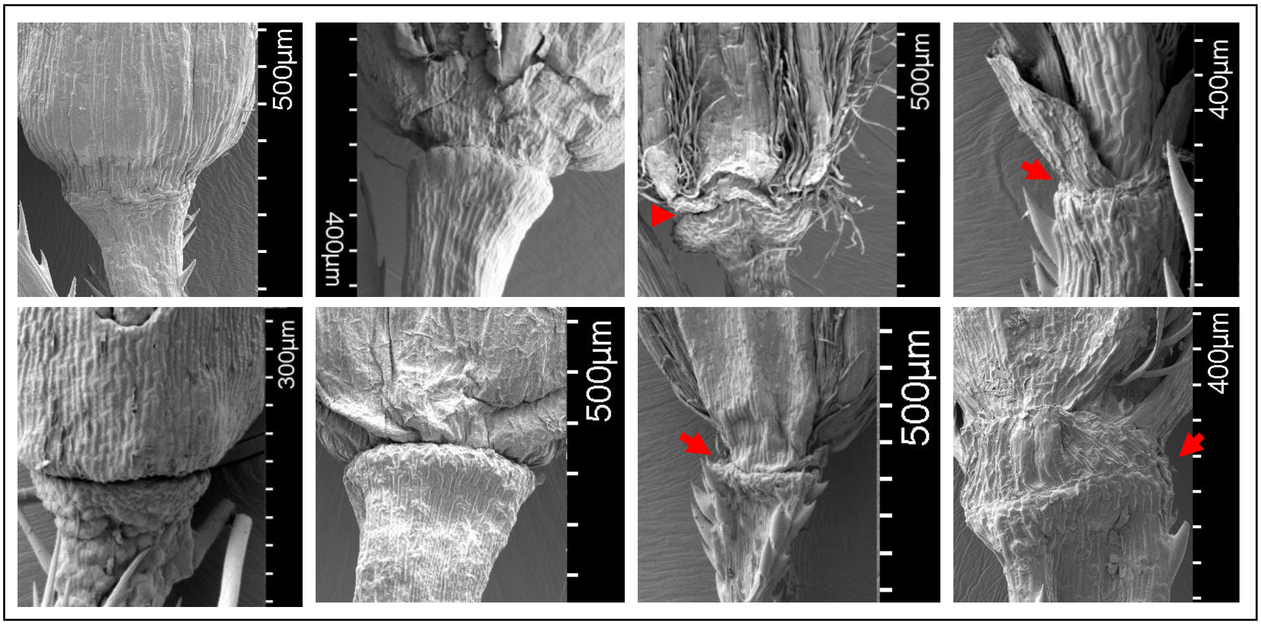


Figure S6 - Scanning Electron Microscope photos of *Digitaria* abscission zones, wild species. Captured at Kew Gardens and the Natural History Museum (UK). Red arrows indicate abscission zone. From top left to bottom right: *D. abyssinica* (Schimper 82), *D. argyrotricha* (Scholz, H. 1096), *D. compressa* (A. Blair Rains 207), *D. debilis* (X.M. van der Burgt 1771), *D. diagonalis* (I. Darbyshire 471), *D. gayana* (F.F. Merklinger 15), *D. perrotettii* (F.F. Merklinger 104), and *D. setigera* (Clayton 5338).

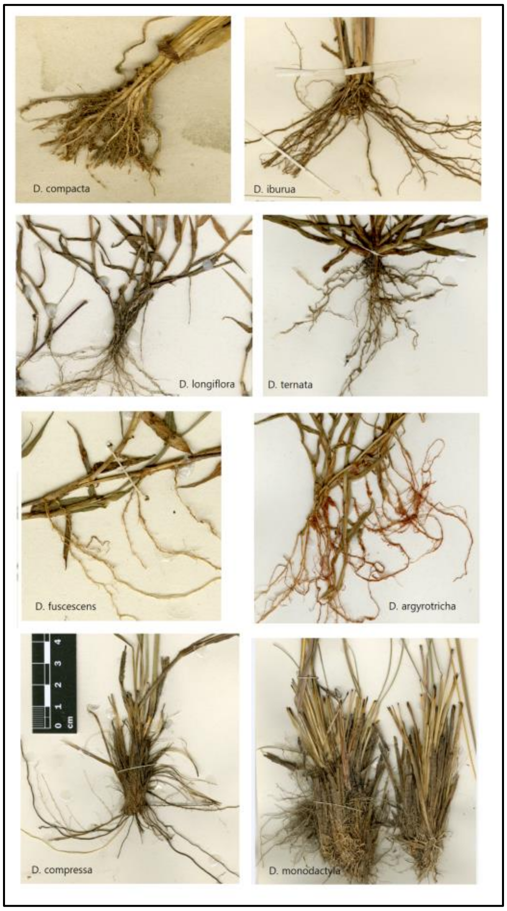


Figure S7 – Growth form morphologies of *Digitaria* species, for cultivated crops (row 1), wild relatives (rows 2 and 3), and early diverging fire-adapted species (row 4). Photos are to same scale, using scale bar in bottom left image.
